# A transcriptomic axis predicts state modulation of cortical interneurons

**DOI:** 10.1101/2021.10.24.465600

**Authors:** Stephane Bugeon, Joshua Duffield, Mario Dipoppa, Anne Ritoux, Isabelle Prankerd, Dimitris Nicolout-sopoulos, David Orme, Maxwell Shinn, Han Peng, Hamish Forrest, Aiste Viduolyte, Charu Bai Reddy, Yoh Isogai, Matteo Carandini, Kenneth D. Harris

## Abstract

Transcriptomics has revealed the exquisite diversity of cortical inhibitory neurons^1–7^, but it is not known whether these fine molecular subtypes have correspondingly diverse activity patterns in the living brain. Here, we show that inhibitory subtypes in primary visual cortex (V1) have diverse correlates with brain state, but that this diversity is organized by a single factor: position along their main axis of transcriptomic variation. We combined *in vivo* 2-photon calcium imaging of mouse V1 with a novel transcriptomic method to identify mRNAs for 72 selected genes in *ex vivo* slices. We used transcriptomic clusters (t-types)^4^ to classify inhibitory neurons imaged in layers 1-3 using a three-level hierarchy of 5 Families, 11 Classes, and 35 t-types. Visual responses differed significantly only across Families, but modulation by brain state differed at all three hierarchical levels. Nevertheless, this diversity could be predicted from the first transcriptomic principal component, which predicted a cell type’s brain state modulation and correlations with simultaneously recorded cells. Inhibitory t-types with narrower spikes, lower input resistance, weaker adaptation, and less axon in layer 1 as determined *in vitro*^8^ fired more in resting, oscillatory brain states. Transcriptomic types with the opposite properties fired more during arousal. The former cells had more inhibitory cholinergic receptors, and the latter more excitatory receptors. Thus, despite the diversity of V1 inhibitory neurons, a simple principle determines how their joint activity shapes state-dependent cortical processing.

---

The cerebral cortex contains a rich diversity of neurons, particularly amongst inhibitory cells. While this diversity was visible to early anatomists^9–11^, the underlying complexity of cortical inhibitory cell types has emerged only with the advent of transcriptomics^1–7^. Single-cell RNA sequencing and Patch-seq analysis suggest that inhibitory neurons of primary visual cortex (V1) are divided into five major Families, named Pvalb, Sst, Lamp5, Vip, and Sncg^3,4,8^. However much finer transcriptomic distinctions exist within these families, with cluster analysis defining 60 different fine inhibitory transcriptomic types (t-types). Moreover, this analysis may underestimate the diversity of cortical inhibitory neurons, which exhibit not only discrete classes but also variations along transcriptomic continua^2,12,13^, which can predict intrinsic physiological properties^2^.

A key open question is whether this fine molecular diversity of cortical inhibitory neurons is mirrored *in vivo* by diverse activity patterns, and whether there are simplifying principles that can help understand the relationship between gene expression and activity in these myriad cell types. Three main methods have been used to characterize the *in vivo* activity of molecularly identified cells. The first is to record from them juxtacellularly and then apply post-hoc morphological reconstruction and immunohistochemistry^14^. This method however has limited throughput, as juxtacellular electrodes can only record one cell at a time. The second is to record from transgenic mice with electrophysiology or 2-photon calcium imaging^15–34^. However, transgenic lines can only identify one molecular group of cells at a time, and the groups these mouse lines identify are broad, containing cells of multiple t-types or even Families. The third and potentially most powerful method is to combine two-photon calcium imaging with ex-vivo molecular identification of the recorded neurons^35–40^. This method can record the activity of large numbers of neurons from multiple cell types simultaneously, and its ability to assign cells to fine molecular types is limited only by the molecular methods used to subsequently identify the neurons.

Here, we used two-photon microscopy to record from large populations of neurons in mouse primary visual cortex (V1), and applied *in situ* transcriptomics to the imaged tissue to localize mRNAs for 72 genes chosen to identify fine inhibitory t-types. While most differences in sensory tuning appeared at the level of main Families (Pvalb/Sst/Vip/Sncg/Lamp5), fine t-types showed significant differences in their modulation by cortical state. These differences in state modulation could be explained in large part by a single genetic continuum, which also correlated with the intrinsic membrane properties and morphology of these t-types as assessed *in vitro^8^,* and with their expression of excitatory and inhibitory cholinergic receptors.

## Identifying inhibitory t-types recorded in vivo

We performed 2-photon calcium imaging in mice expressing mCherry in inhibitory neurons (Gad2-T2a-NLS-mCherry), injected with a pan-neuronal GCaMP6m virus (AAV1-Syn-GCaMP6m-WPRESV40), and then applied *in situ* transcriptomics to sagittal slices of the imaged region. The expression of mCherry allowed these neurons to be localized *in vivo* and *ex-vivo* and thus to serve as fiducial markers for registration of the two imaging modalities. During imaging, mice were free to run on an air-suspended Styrofoam ball, and their behavioural state was monitored through facial videography. Spontaneous activity was recorded in front of a blank screen, and visual responses were elicited by presenting stimuli such as drifting gratings and natural images. Recordings typically spanned 0-250 μm below the brain surface, targeting cortical layers 1-3. At the end of each session, we obtained a high-resolution 2-photon z-stack volume (Fig. 1a). After functional imaging was complete, brains were removed and frozen unfixed, and the imaged volume was cut into 15 μm thick sections with a cryotome.

**Figure 1.**
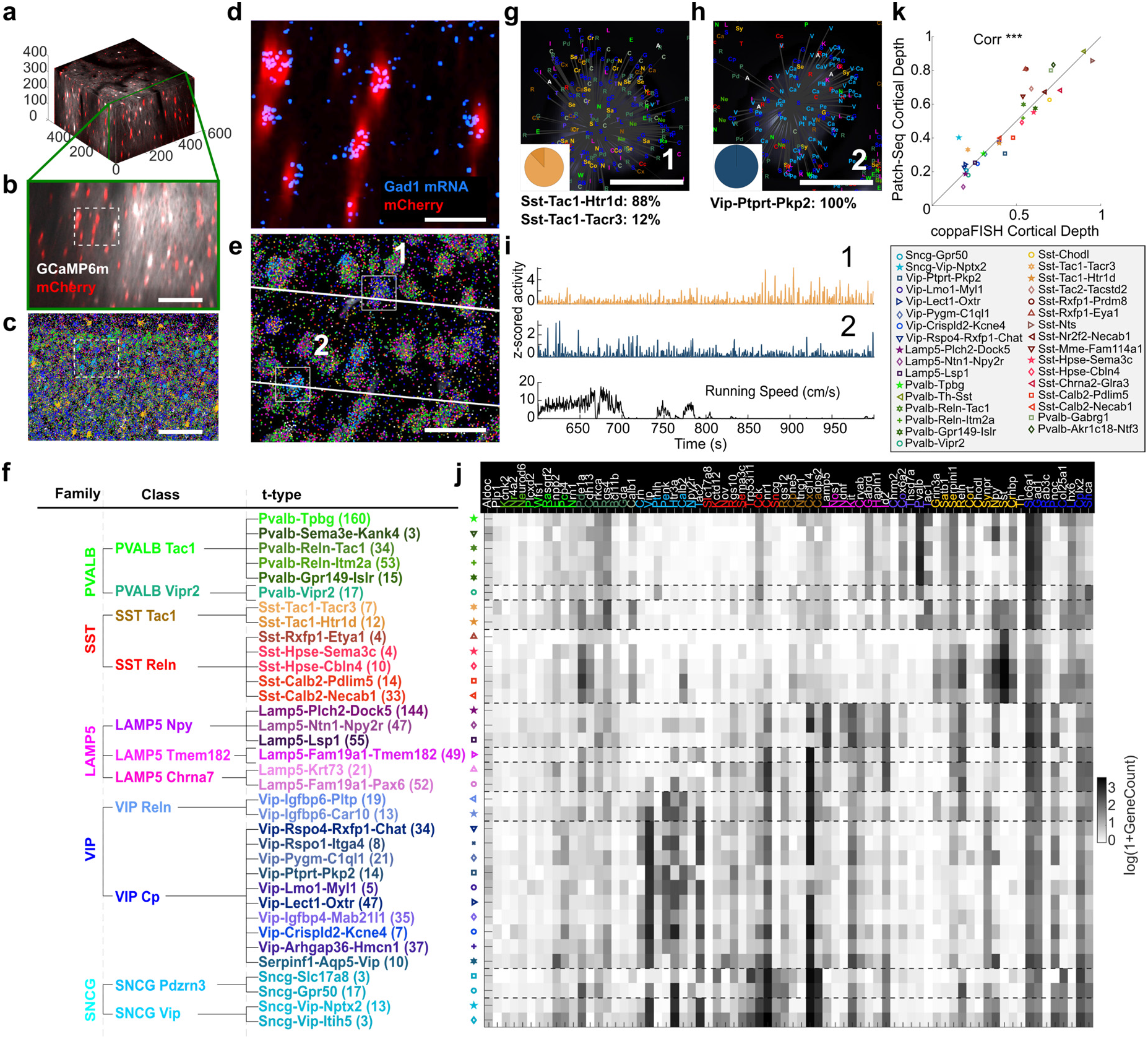
Post-hoc transcriptomic identification of recorded neurons. **a**, 3D representation of an example reference Z-Stack (white: GCaMP6m, expressed virally in all neurons, red: mCherry expressed transgenically in inhibitory neurons). Axis scale in μm. **b**, Digital sagittal section of this Z-Stack (maximum intensity projection of a 15 μm slice), same colours as in **a**. Scale bar: 100 μm. **c**, Portion of the *ex vivo* section aligned to the cut in **b** after 72-fold mRNA detection with coppaFISH. Individual dots represent detected mRNAs (see top of **j** for colour code). Scale bar: 100 μm. **d**, Expanded view of the dashed rectangle in **b** and **c** showing *in vivo* mCherry fluorescence (red) and *ex-vivo* Gad1 mRNA detections (blue). Scale bar: 20 μm. **e**, Expression of all 72 genes in this same region, plotted as in **c**. White lines indicate two functional imaging planes. Gray background: DAPI stain for cell nuclei. Scale bar: 20 μm. **f**, Hierarchical classification of recorded cell types into 5 Families, 11 Classes, and 35 t-types. Number of unique cells of each t-type given in parentheses. **g**, **h**, Higher magnification view of Cells 1 and 2 from **e**. Gene detections are indicated by coloured letters (code at top of **j**). Gray background: DAPI image. Insets: pie plots indicating the posterior probabilities of assignment to different t-types. Scale bar: 5 μm. **i**, Deconvolved calcium traces for the two example cells, shown together with running speed. **j**, Mean expression of the 72 detected genes (pseudocoloured as log(1 +GeneCount)) for the 35 t-types ordered as in **f** (n=4 animals). **k**, Comparison between the mean cortical depth of each t-type found using coppaFISH (on N = 14 sections from a brain in which mRNAs were detected down to Layer 6), and the cortical depth found by an independent study using Patch-seq^8^. (r=0.91, p<0.001). Only t-types with at least 3 cells for each dataset were considered. The black line represents equality.

To identify the locations of 72 pre-selected genes we developed a method termed coppaFISH (combinatorial padlock-probe-amplified fluorescence *in situ* hybridization), which is a development of a previous approach of *in situ* sequencing^41^. This method amplifies selected transcripts *in situ* using barcoded padlock probes^42–45^ and reads out their barcodes combinatorially through 7 rounds of 7-colour fluorescence imaging (Methods; Extended Data Fig. 1). The method detected 148±59 transcripts per cell (mean ± SD). The slices were aligned to the *in vivo* z-stacks with a pointcloud registration algorithm, using inhibitory neurons identified *in vivo* with mCherry and *ex vivo* through gene expression, as fiducial markers (Fig. 1b-e; Extended Data Fig. 2). We applied this method to 17 recording sessions from 4 mice, obtaining 89±30 (mean ± SD) molecularly-identified inhibitory cells together with 393±173 pyramidal neurons per session, making a total of 1,028 unique molecularly identified inhibitory cells (some of which were recorded in multiple sessions; Supplementary Data File 1).

We classified these inhibitory cells using a 3-level hierarchy (Fig. 1f). The lowest hierarchical level (“t-type”) comprised the fine transcriptomic clusters defined by Tasic et al.^4^, and the top level (“Family”) was the Pvalb, Sst, Lamp5, Vip, and Sncg groupings defined by these same authors. An intermediate level (“Class”) was suggested by UMAP analysis of scRNA-seq data (Extended Data Fig. 3), which revealed collections of t-types that we could putatively associate to morphological cell types (see Methods for full explanation). We named these intermediate-level Classes Pvalb-Tac1 (putative Pvalb-basket cells), Pvalb-Vipr2 (chandelier cells), Sst-Reln (Martinotti cells); Sst-Tac1 (non-Martinotti Sst cells); Lamp5-Npy (neurogliaform cells); Lamp5-Tmem182 (canopy cells); Lamp5-Chrna7 (layer 1 alpha7 cells); Vip-Reln (layer 1 Vip cells); Vip-Cp (other Vip cells); Sncg-Pdzn3 (Large Cck cells); and Sncg-Vip (Small Cck/Vip cells). UMAP analysis (Extended Data Fig. 3) suggested that while Classes were usually discrete, their constituent t-types often merged continuously into each other, tiling dimensions of continuous variability of inhibitory neurons.

Cells functionally imaged *in vivo* were assigned to a t-type (and thus also a Class and Family) using pciSeq^41^, a Bayesian algorithm that computes for each cell a probability distribution over cell types defined by previous scRNA-seq data. Expression levels were sufficient to assign cells with high probability to a single t-type (Fig. 1g-i, Extended Data Fig. 4), and we therefore assigned each cell to a single t-type of maximum *a posteriori* probability. As expected from the restriction of 2-photon imaging to the superficial layers, the imaged cells were assigned to just 35 of the 60 total t-types defined by the original scRNA-seq study. The number of cells recorded varied across types (Fig. 1f), and t-types to which less than 3 cells were identified were excluded from further analysis (eight cells in total). The gene expression for the 72 genes in our panel showed consistent differences across the 35 t-types recorded (Fig. 1j).

To verify the accuracy of our cell type assignments, we performed two analyses using independent data. First, we took advantage of the fact that different fine inhibitory t-types reside at different depths in V1, as revealed by a recent Patch-seq study^8^. We found that the depth distribution of the t-types assigned by our method closely matches that found by this independent study (Fig. 1k). Importantly, this did not only reflect depth differences between the main inhibitory Families (p<0.001, ANCOVA controlling for Family) or even Classes (p<0.001, ANCOVA controlling for Class). For example, while Sst-expressing neurons are most often found in deep layers, specific t-types such as Sst-Calb2-Necab1 were localized in superficial layers by both our method and the independent Patch-seq data. Second, we compared the functional recordings to 2-photon calcium imaging that identified cells with three transgenic lines (Sst, Pvalb, Vip)^19^. Analysing our data after grouping together cells expected to be labelled in each of these lines, we found results consistent with previous work (Extended Data Fig. 5). We thus conclude that our methods accurately identify fine t-types, and that the functional correlates of these cells match those previously observed at the Family level with previous methods.

## State modulation of inhibitory t-types

We next asked to what extent fine transcriptomic types affect a neuron’s *in vivo* activity patterns. We generated raster plots showing the simultaneous activity of V1 populations, with all inhibitory neurons identified to fine transcriptomic types (Fig. 2a). Examining these rasters during spontaneous activity revealed complex patterns of correlated activity, which varied with ongoing behaviour as measured by two assays of arousal: locomotion and pupil diameter. It has been possible to plot such rasters before, but without identifying the transcriptomic identity of the recorded neurons. We therefore first asked how activity of the identified cell types depended on cortical state, a correlate of these assays of arousal^21,29,32,33,46–48^.

**Figure 2.**
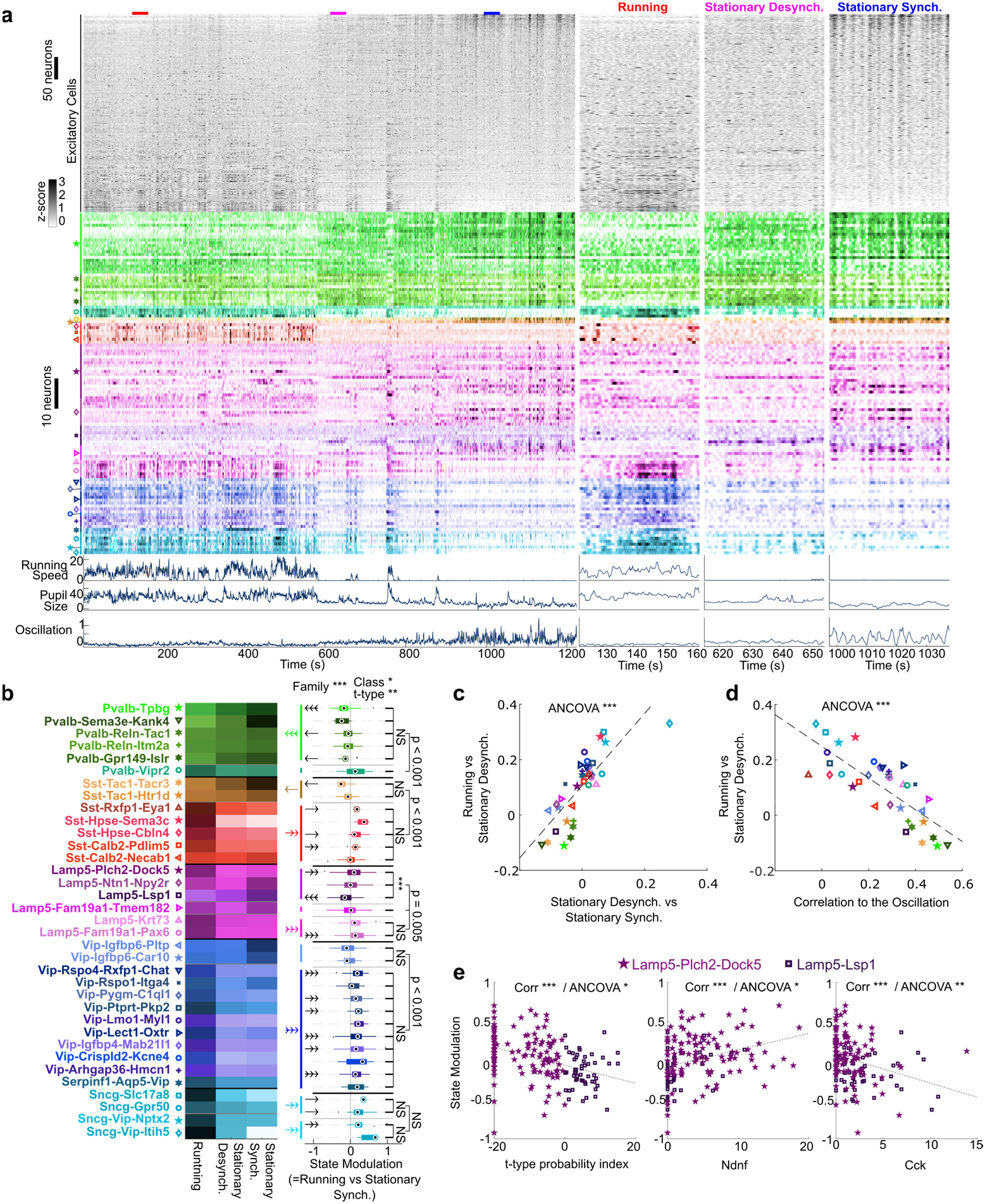
State modulation of inhibitory t-types. **a**, Raster of spontaneous neuronal activity (blank screen). Each row represents a neuron. Top, excitatory cells (ECs; mCherry-negative), sorted by weight on the first principal component (PC) of their activity. Middle: inhibitory cells, grouped and coloured by t-type (symbol code in **b**). Bottom: running speed (cm/s), pupil size (pixels), and mean activity of the 10% of ECs with most negative PC weights. Three columns on right: expanded view of time windows marked in colour above main raster, illustrating three behavioural states. **b**, Left: pseudocolour representation of mean activity of each t-type in each state. Right: distribution of state modulation (Running vs. Stationary Synchronized) for cells of each t-type (n=4 animals; 17 sessions). Top p-values: Omnibus test for Family/Class/t-type effects. p-values on right: post-hoc tests for effect of Class within each Family; stars on right for effect of t-type within each Class (Benjamini-Hochberg corrected). Coloured/black arrows on left: significant state modulation for each Class/t-type (Benjamini-Hochberg corrected, number of arrowheads indicates significance). **c**, State modulation for Running vs. Stationary Desynchronized states, against modulation for Stationary Synchronized vs. Desynchronized states. Each glyph shows median values for a t-type, symbols as in **b** (p<0.001, ANCOVA controlling for session). **d**, Modulation for Running vs. Stationary Desynchronized states against locking to the synchronized-state oscillation. Each glyph shows median values for a t-type. (p<0.001, ANCOVA controlling for session). **e**, State modulation for cells in the Lamp5-Plch2-Dock5 and Lamp-Lsp1 t-types, against t-type probability index (log(pType1/pType2); left), or Ndnf (middle) and Cck (right) gene expression. These three variables correlated significantly with state modulation (p<0.001, Pearson correlation), even controlling for a common effect of t-type (p<0.05, ANCOVA). Dashed lines: linear fits. *, p<0.05, **, p<0.01, ***, p<0.001; 1, 2, or 3-headed arrows in **b** indicate the same significance levels, with direction indicating the sign of the modulation.

We characterized cortical state using the activity of the excitatory population. As previously described^49^, some excitatory cells (positively weighted on the first principal component of population activity) were more active when the mouse was aroused (fast running, large pupil) while other excitatory cells (with negative weights) fired during inactive periods (no running, small pupil). Additionally, we found that behavioural inactivity was sometimes accompanied by low-frequency fluctuations in population activity, which strongly entrained the excitatory neurons as visible in the mean activity of the negatively weighted cells (Fig. 2a). The frequency of these fluctuations cannot be determined from our data as our two-photon microscope’s 4.3 Hz sampling rate aliases frequencies above 2.15 Hz, but they likely correspond to the 3-6 Hz oscillation that has been described in mouse visual cortex and hypothesized as a homolog of the primate alpha rhythm^50–53^. We thus distinguished three cortical states corresponding to decreasing levels of arousal: periods when the mouse is running; periods when the mouse is stationary but the network desynchronized; and stationary periods with synchronized, oscillatory population activity. To quantify a cell’s modulation by cortical state we compared the activity of each cell during the two extreme states: Running vs. Stationary Synchronized.

Statistical analysis of differences between inhibitory cell types must avoid two potential confounds. First, the large number of t-types presents a potential multiple comparisons problem. Second, different recordings will by chance sample different proportions of each cell type, and thus recording-to-recording variability could be mistaken for variability between cell types. To avoid these confounds, we used a hierarchical permutation test (Methods), which tests for a main effect of Family, Class, or t-type on a variable of interest by shuffling the molecular labels of cells within an experiment, at each of the 3 hierarchical levels.

The hierarchical permutation test revealed significant differences in state modulation at all three levels: Family, Class, and t-type (Fig. 2b). Post-hoc tests revealed large differences in activity across Classes in the Pvalb Family: Pvalb-Tac1 cells were strongly active during oscillatory states and less active during running, while Pvalb-Vipr2 cells showed the opposite behaviour (consistent with previous results^54^). Likewise, in the Sst Family, Sst-Tac1 cells were most active during synchronized states while Sst-Reln cells were more active during running. Similar dichotomies were observed in the Lamp5 Family. Vip and Sncg cells were more active during running, except for Vip-Reln cells, which showed the opposite behaviour. Post-hoc statistical tests at the t-type level revealed that the most prominent difference was within the Lamp5-Npy (putative neurogliaform) Class; a trend toward difference was also seen in Sst-Reln (putative Martinotti) cells (p<0.05, significant on its own but not after Benjamini-Hochberg correction). Analysis of the Stationary Desynchronized state revealed intermediate activity compared to the two extreme states (Extended Data Fig. 6a-b). The modulations between either one of the extreme states and the intermediate state (Stationary Desynchronized) were strongly correlated, indicating a linear progression of neural activity across the three states (Fig. 2b-c). A t-type’s state modulation was correlated with its degree of phaselocking to the Synchronized state oscillation, with t-types more active during Running less locked to the oscillation during the Stationary Synchronized periods (Fig. 2d).

The dependence of state modulation on t-type was consistent with smooth variation along a continuum of genetic types, rather than a sharp difference between discrete groups. For example, amongst t-types of the Lamp5-Npy Class, Lamp5-Lsp1 cells were most active in the Synchronized state, while Lamp5-Plch2-Dock5 cells fired more during Running. The division between these t-types however reflects a largely arbitrary dividing line along a continuous dimension of genetic variability (Extended Data Fig. 3). To test if their divergent state modulation followed a continuous, rather than discrete transcriptomic variable, we quantified each imaged cell’s position along the continuum by its ratio of posterior probabilities of assignment to the two t-types. We observed a smooth dependence of state modulation along this continuum, which ANCOVA analysis showed depended on this continuous genetic score better than on discrete t-type assignment (Fig. 2e). Similar continuous dependence was visible at the single-gene level, with state modulation within Lamp5-Npy cells correlating with expression of Cck and Ndnf (Fig. 2e) even after controlling for t-type. Similar results were seen for Sst-Reln t-types (Extended Data Fig. 6c).

## Sensory responses of inhibitory t-types

We next probed the responses of different inhibitory types to visual stimuli: drifting gratings of various sizes and orientations, and natural images. Unlike state modulation, visual responses showed significant differences only at the level of Families, not Classes or t-types.

Most inhibitory cell types contained neurons responding to grating stimuli (Fig. 3a-d; Extended Data Fig.7). Pvalb and Sst cells responding to gratings were almost exclusively excited by them, whereas Sncg cells, whose visual responses to our knowledge have not yet been studied, were almost exclusively inhibited. Lamp5 and Vip cells contained a mixture of excited and inhibited cells, with Vip cells more often excited. Orientation and direction tuning was relatively low for most Families^22,28,36,55–57^, with a slight tendency for Sst and Vip cells to show stronger tuning. Most cells showed significant coding of natural image stimuli, which again differed significantly between Families, being weakest for Sncg cells.

**Figure 3.**
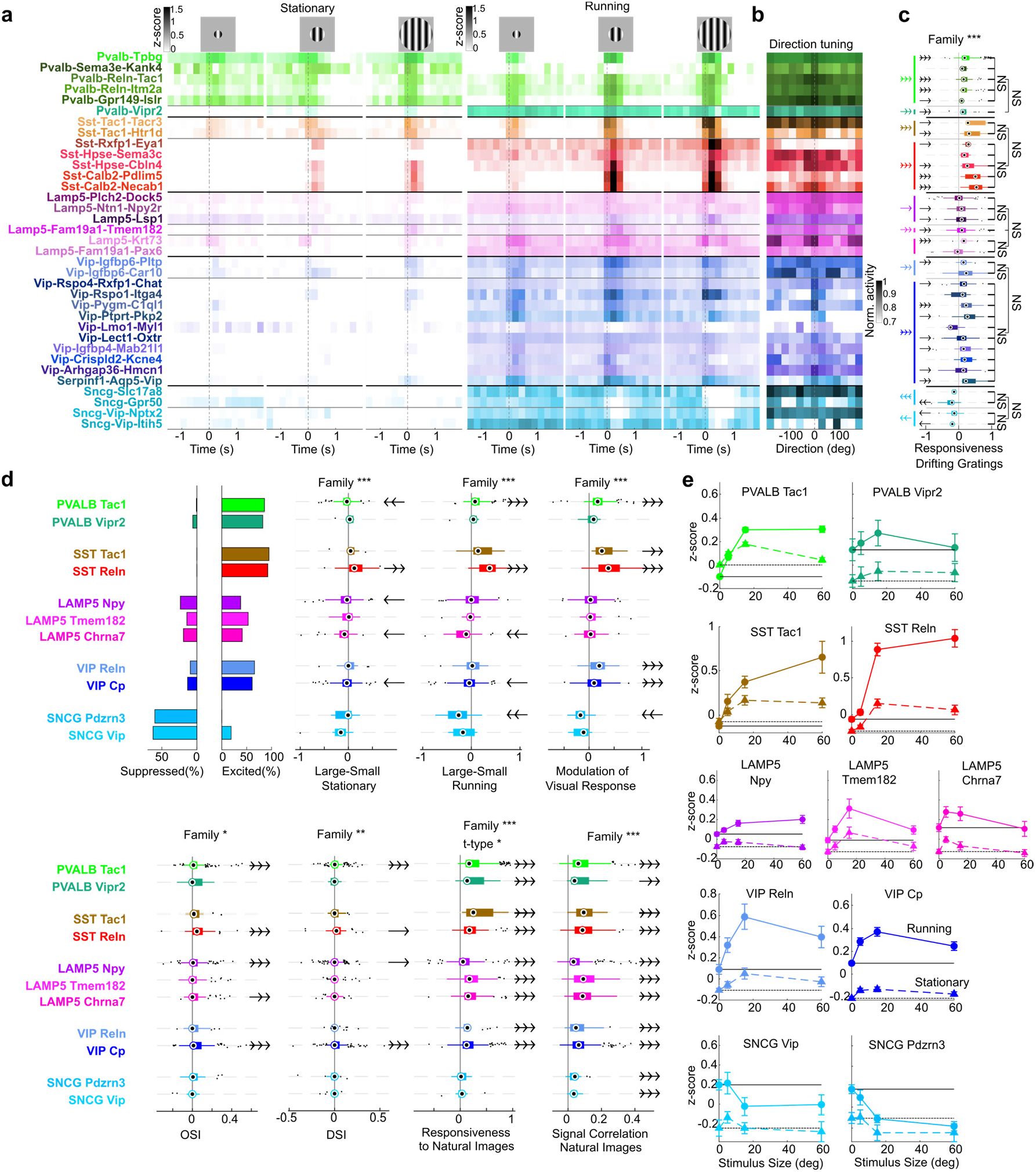
Sensory responses of inhibitory t-types. **a**, Pseudocolour rasters of trial-averaged activity around onset of drifting grating stimuli (duration 0.5s), for different stimulus sizes (5°, 15°, 60°) and locomotor states. Each row shows average activity of a t-type. Dashed grey lines: stimulus onset. **b**, Cross-validated direction tuning curves for each t-type, shown in pseudocolour as a function of grating direction. Tuning curves were averaged over odd trials, shifted and normalized according to the preferred direction on even trials, and averaged across cells of the same t-type. **c**, Hierarchical analysis of responsiveness to drifting gratings (measured at the stimulus size eliciting the largest negative or positive response), plotted as in Fig. 2b. **d**, Additional statistical analyses of visual stimulus responses. Top row, from left: fraction of cells of each Class significantly excited or suppressed by grating stimuli; hierarchical analysis of differences between mean response to large and small gratings in stationary and running conditions, and modulation of visual response by running averaged over all sizes, plotted as in Fig. 2b but only showing the Class level. Bottom row: hierarchical analysis of orientation and direction selectivity indices, responsiveness and tuning to natural image stimuli. **e**, Size tuning curves for each Class. Dashed lines and triangular markers correspond to mean stimulus response during stationary periods. Solid lines and circular markers correspond to mean stimulus response during running epochs. Black dashed and solid lines indicate respectively the average activity during baseline for Running or Stationary (interstimulus periods).

The most striking difference in the grating responses of different inhibitory cell types was in their tuning to grating size and its modulation by cortical state (Fig. 3e). Size tuning was significantly modulated at the Family level: as previously reported^16,19^, Sst cells showed little or no surround suppression, with strong responses to large stimuli. Sncg cells showed a strikingly opposite pattern in which they were progressively more suppressed by larger stimuli (Fig. 3e). Modulation of grating response by locomotion was significantly different between Families, with Sst, Pvalb, and Vip cells showing various degrees of increase with locomotion and Sncg a decrease.

In summary, sensory responses showed significant differences between Families, but not between Classes and t-types. The most striking differences between Families were in size tuning and its modulation by state. A lack of statistical significance of course does not exclude the possibility that t-types may differ in sensory tuning in ways too small for our methods to detect; but the fact that the same statistical tests found t-type differences in state modulation suggests that any such differences in sensory tuning are likely to be subtle.

## A single genetic axis predicts state modulation

Although the number of inhibitory t-types is large and their state modulation is diverse, we found that a large portion of this diversity can be explained by a single genetic axis. This axis was defined independently of the physiological data: we simply computed the first principal component of the gene expression vectors measured *in situ* (genetic PC 1, or gPC1). A similar approach previously applied to scRNA-seq data from CA1 inhibitory neurons revealed a continuum ranging from cells targeting excitatory cell somas at one end, to cells targeting distal dendrites and inhibitory neurons at the other^13^. Applying genetic PCA to the *in situ* transcriptome of our cells revealed a similar continuum (Fig. 4a). The classes with the most negative gPC1 values were Pvalb-Tac1 and Sst-Tac1 cells; those with the highest were Sncg, Vip, Lamp5-Chrna7 and Lamp5-Tmem182 cells; Sst-Reln and Lamp5-Npy Classes occupied the centre of the continuum. Genes negatively correlated with the continuum included Gad1 and Slc6a1 (Fig. 4a), which are involved in GABA synthesis and transport, consistent with previous analysis suggesting that cell types negatively weighted on this continuum exert stronger inhibition on their targets and have faster metabolic rates^13^.

**Figure 4.**
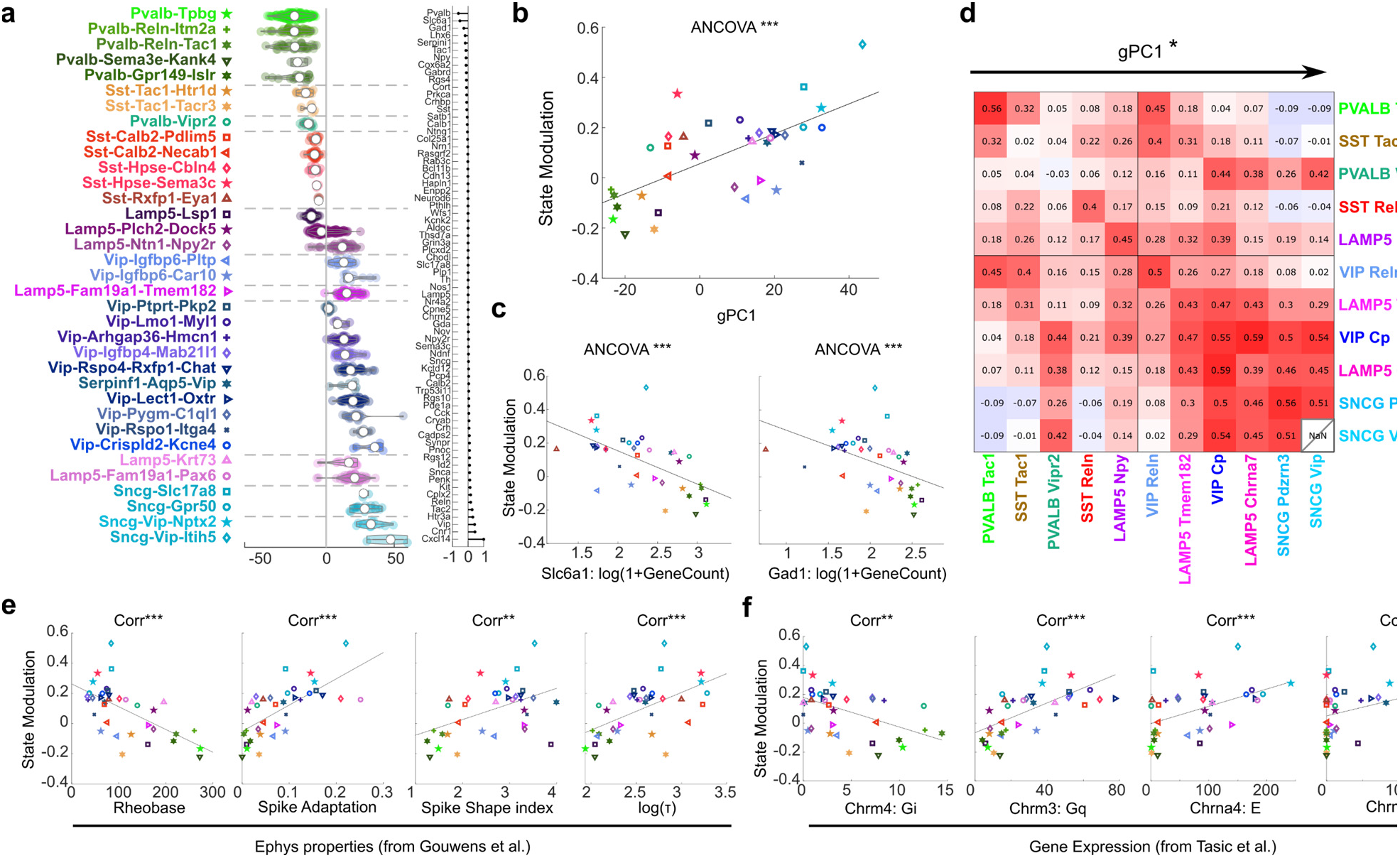
A single genetic axis explains state modulation. **a**, Left: violin plots showing distribution of the first genetic principal component (gPC1) for each t-type. Right: weighting of each gene in this principal component. **b**, Correlation between state modulation and gPC1. Each glyph represents median values for a t-type, symbols as in **a** (p<0.001, ANCOVA controlling for session). **c**, Correlation between state modulation and expression of Slc6a1 and Gad1 measured *in situ.* Each dot represents median values for a t-type, coded as in **a.** (Slc6a1: p<0.001, ANCOVA controlling for session; Gad1: p<0.001, ANCOVA controlling for session). **d**, Matrix of pairwise correlations between simultaneously recorded Classes. The Classes are sorted by gPC1, showing a significant effect of gPC1 on the pairwise correlations (p=0.013, permutation test). **e**, Correlation between state modulation and electrophysiological properties measured by an independent Patch-seq study^8^. Each symbol represents median values for a t-type, coded as in **a**. Rheobase: r=0.39; Spike Adaptation: r=0.47; Spike Shape Index: r=0.23; log(τ): r=0.32. (Significance: Pearson correlation). **f**, Correlation between state modulation and Cholinergic receptor expression obtained from an independent scRNA-seq study^4^. Each symbol represents median values for a given t-type, coded as before. Chrm4: r=0.24; Chrm3: r=0.38; Chrna4: r=0.23; Chrna5: r=0.12. Gq and Gi indicate metabotropic receptors coupled to a Gq (excitatory) and Gi (inhibitory) pathways, E indicates excitatory ionotropic receptor. Correlations of state modulation with excitatory cholinergic receptor expression were higher than with inhibitory receptor expression (including those not shown here; p = 0.01, ANOVA; only receptors with > 2 counts in at least 10 t-types were considered, making 10 in total). *, p<0.05, **, p<0.01, ***, p<0.001; Dashed lines are linear regression fits.

The state modulation of a t-type correlated with its position along the genetic continuum gPC1 (Fig. 4b). Cells with negative gPC1, such as Pvalb-Tpbg (putative basket cells) were most strongly active in Synchronized states, while cells with positive gPC1 such as Sncg cells were most active during Desynchronized and Running states. State modulation was significantly correlated with gPC1, (p<0.001, ANCOVA controlling for session; Fig. 4b). For example, Sst-Tac1 cells, which show a faster-spiking physiological profile than Sst-Reln cells^8^, had the lowest gPC1 values and greatest preference for oscillatory states amongst the Sst population (Fig. 4b). These effects could be seen at a single-gene level, with a t-type’s state modulation negatively correlated with expression of Slc6a1 and Gad1 (p<0.001, ANCOVA controlling for session). Thus, different inhibitory t-types have diverse relationships to cortical state, but these relationships can be at least partially predicted by a single genetic axis, with the side of this axis associated with stronger GABA synthesis and release showing more activity in oscillatory states.

The main gPC1 axis also largely predicted correlations between the spontaneous activity of inhibitory Classes, with positive correlations between Classes of similar gPC1 values, and negative correlations between Classes of opposite gPC1 values (p<0.05, per-mutation test; Fig. 4d). This also held true when considering correlations computed within any of the three states independently (Extended Data Fig. 8).

A cell type’s state modulation and position on the gPC1 axis also correlated with many aspects of its intrinsic physiology and morphology (Fig. 4e). To demonstrate this, we analysed data from an independent Patch-seq study^8^. Transcriptomic types that were active during synchronized states (low arousal levels) had faster membrane time constants and spike repolarization speeds, more hyperpolarized resting potential, lower membrane resistance, larger rheobase (i.e. the amount of current required to drive spiking), and less spike frequency adaptation (Fig. 4e; Extended Data Fig. 9a). Transcriptomics types active during running had the opposite properties. This Patch-seq data also revealed an intriguing correlate of gPC1 and axonal morphology. Within the Sst and Lamp5 Families, cells with larger values of gPC1 (which thus would show more activity in alert states *in vivo*) had a greater fraction of their axon in layer 1, and a smaller fraction in layer 2/3 (p<.001, Pearson correlation with Benajamini-Hochberg correction; Extended Data Fig. 9b). This correlation was not seen for the other Families, for which axonal projections to layer 1 were rare.

Finally, we hypothesized that variation in state modulation along the gPC1 axis might reflect variation in cholinergic receptor expression. Acetylcholine levels are higher in active states and contribute to cortical desynchronization^58–64^. Moreover, acetylcholine differentially affects inhibitory neuronal types by acting through different receptors, with nicotinic and G_q_-coupled muscarinic receptors exciting some inhibitory types and G_i_-coupled muscarinic receptors inhibiting others^65–70^. To test this hypothesis, we examined the correlation between state modulation and expression of each cholinergic receptor type across t-types. Consistent with the hypothesis, we found positive correlations between state modulation and the expression level of all nicotinic or G_q_-coupled muscarinic receptors, and negative correlations between state modulation and expression levels of G_i_-coupled muscarinic receptors (Fig. 4f; excitatory receptors significantly more positively correlated than inhibitory receptors, p<0.05, ANOVA). This suggests that differential expression of cholinergic receptor subtypes may contribute to the smooth variation of state modulation along the main axis of genetic variation gPC1.

## Discussion

By genetically identifying the transcriptomic types of simultaneously recorded V1 neurons, we discovered fine functional differences across cellular t-types and a simple ordering along a main axis of genetic variation. These differences and this ordering were seen not in the sensory responses of the neurons – which differed primarily across high-level Families – but rather in the relation of their activity with cortical and behavioural state. State modulation can vary significantly between fine t-types within a Class, but this appears to reflect continuous genetic variation rather than discrete t-types. Furthermore, a single, simple, axis of genetic variation across inhibitory cells – the first principal component of gene expression (gPC1) – largely explains differences in state modulation between t-types, and predicts their spontaneous correlations. This genetic axis also correlates with a t-type’s membrane physiology, layer 1 axon content, and expression of excitatory and inhibitory cholinergic receptors.

The diversity that we observed across t-types may explain why previous reports of state modulation of different interneuron types, based on transgenic lines, have at times given apparently conflicting results. Previous work has uniformly shown that the activity of Vip-Cre labelled neurons in V1 is enhanced by running^19,21,29,33,71^ and our data are fully consistent with this. Recent work has shown that most V1 cells labelled in Ndnf-Cre mice fire more in aroused states^72^.

This is again consistent with our results: Ndnf is found in the Lamp5 family but not the Lamp5-Lsp1 t-type, which was the only Lamp5 t-type we found to have significantly negative state modulation. Measurements of running modulation in Pvalb-Cre mice (which will label mainly basket cells in V1) have shown mixed results^19,21,29^, which may be explained by an effect of cell depth on running modulation: cells above 300 μm show primarily suppression by running and cells below that depth primarily excitation^19^. Our data involved only cells above 300 μm depth and showed close to uniform negative state modulation in Pvalb-Tac1 (putative basket) cells but positive modulation in Pvalb-Vipr2 (putative chandelier), consistent with recent data from Vipr2-Cre mice^54^. We speculate that the deeper-layer cells positively modulated in Pvalb-Cre mice correspond to additional Pvalb-Tac1 t-types not recorded in this study. An additional factor that may explain differing results in previous work is light level. Work in Sst-Cre mice has shown that running suppresses activity in pitch darkness, but has mixed effects in light^19,21,29,32,72^; our data were conducted in light, and we speculate that the mixed effects seen in Sst-Cre mice might reflect a Class difference, with Sst-Tac1 cells suppressed but Sst-Reln cells activated.

It is remarkable that a single transcriptomic dimension – derived from gene expression patterns without regard to physiological properties – correlates with state modulation that we measured *in vivo*, with intrinsic physiology measured *in vitro^8^,* and with the expression of cholinergic receptors with opposite signs for excitatory and inhibitory receptors^4^. The continuum we observed is similar to one previously described in scRNA-seq data from CA1 inhibitory neurons^13^, but with one notable exception: in CA1, Sncg t-types occupied multiple locations along the continuum, rather than all being at the positive end as in V1. This might be related to the existence of fast-spiking CCK basket cell subtypes in CA1^73^, and the fact that CA1 Sncg cells can be inhibited by locomotion^20^.

Although we have focused here on one dimension of arousal/desynchronization, the space of cortical states is unlikely to be one dimensional, as multiple dimensions of V1 excitatory cell activity correlate with ongoing behaviour^49^. Characterizing how the multidimensional space of cortical states relates to multiple inhibitory classes and ongoing behaviours remains an important topic for future work, as does understanding whether this relationship varies between different cortical regions. High-throughput application of the current methods will make this possible.

The existence of these correlations with gPC1 suggests that many observations, made on individual inhibitory types, could be consequences of a general principle applying to all interneurons. For example, acetylcholine has been shown to have diverse effects on different inhibitory types^65,67–69^, such as the classical “cholinergic switch”^70^ whereby fast spiking (putative Pvalb basket) cortical neurons are inhibited by muscarinic receptors but low-threshold spiking (putative Sst Martinotti) neurons are excited by nicotinic receptors. This result is consistent with the receptor expression profile of these Classes, and with our finding that the Pvalb-Tac1 Class is inhibited, and the Sst-Reln Class excited in Desynchronized and Running states. In fact, our data suggest that the behaviour of these two cell Classes is a manifestation of a more general principle: at least in superficial V1, inhibitory cells with lower gPC1 values exhibit physiological properties closer to Pvalb basket cells, lower levels of nicotinic and excitatory muscarinic receptors, more inhibitory muscarinic receptors, and negative state modulation, and the reverse is true for cells with larger gPC1 values. Differences in cholinergic receptor expression likely contribute to differences in *in vivo* state modulation: acetylcholine levels are largest in locomotion and lowest in synchronized states, and state modulation of at least some interneuron classes depends on cell-type-specific nicotinic and muscarinic currents^21,69^. Direct cholinergic input is of course unlikely to be the only factor mediating state dependence of an interneuron class: interneurons receive input from pyramidal cells, and from each other in specific ways such as the well-known “disinhibitory circuit”^21,25,69^. Nevertheless, the correlation of cholinergic receptor expression and state modulation we observed suggests that cell-type-specific cholinergic modulation may play a substantial role, at least in superficial V1.

What computational role might be served by this state-dependent switch in the activity of different inhibitory cell types? Our data are consistent with a long-standing view that alert states and cholinergic modulation biases cortex towards feedforward inputs from primary thalamus, and away from top-down inputs from elsewhere in cortex^74–78^ (Fig. 5). Indeed, the Classes most suppressed by alert states (putative Pvalb basket and Sst non-Martinotti) preferentially target thalamorecipient layers 4 and 5b, while the Sncg, Lamp5, Sst-Martinotti and Vip cells more excited in alert states preferentially target either interneurons, or pyramidal cells in other layers^79–81^. Our data furthermore suggests that the degree of state modulation for Sst and Lamp5 neurons correlates with their axonal innervation of layer 1, which receives top-down input. Opposing cholinergic modulation of these inhibitory classes might thus alter the balance between bottom-up and top-down inputs.

**Figure 5.**
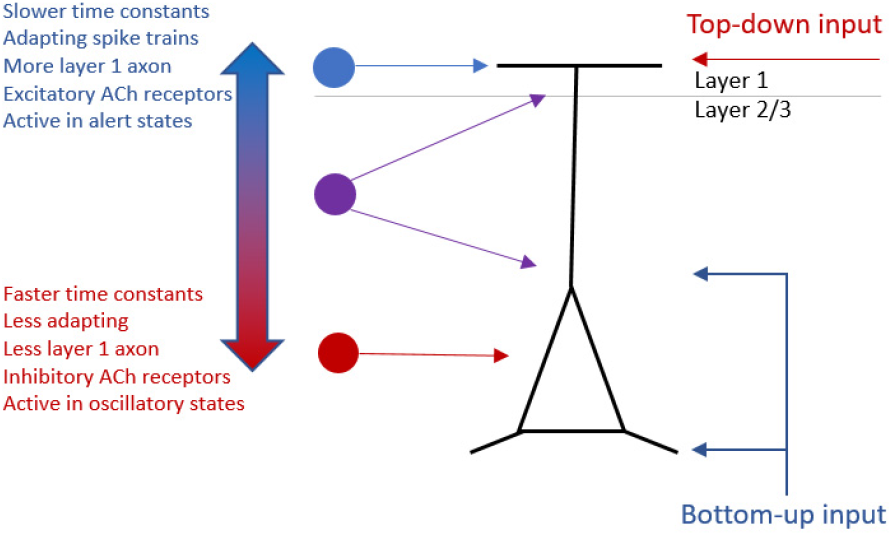
Model for how state-dependent modulation of inhibitory subtypes could bias cortex towards bottom-up or top-down inputs.

In summary, while V1 inhibitory neurons are very genetically diverse, we found that their sensory tuning is determined largely by their top-level transcriptomic Family, and their state modulation can be predicted in large part from a single genetic axis that also correlates with their intrinsic physiology, morphology, and cholinergic receptor expression. As emerging experimental techniques allow for ever-greater amounts of information to be collected on the physiology, connectivity, and firing correlates of cortical interneuron classes, these simple principles may help organize this information.

## Methods

All experimental procedures were conducted in accordance with the UK Animals (Scientific Procedures Act) 1986. Experiments were performed at University College London under personal and project licences released by the Home Office following appropriate ethics review.

### Mice

Experiments were performed on mice aged between 12 and 15 weeks maintained on a 12-h light/dark cycle, at 20–24 °C and 45–65% humidity, in individually ventilated cages. For post-hoc identification of transcriptomic t-types, four (two males and two females) Gad2-T2a-NLS-mCherry transgenic mice (Stock No: 023140, The Jackson Laboratory), expressing the red fluorescent protein mCherry in the nuclei of Gad2 expressing cells, were used. For comparison to transgenic mouse lines (Extended Data Fig. 5), additional experiments were performed as in Ref.^19^ using one male Pvalb<tm1(cre)Arbr> and 2 male, 1 female Sst<tm2.1(cre)Zjh> crossed with Gt(ROSA)26Sor<tm14(CAG-tdTomato)Hze>.

### Surgical procedures

On the day of surgery, mice were anaesthetized with isoflurane (1–2% in oxygen), their body temperature was monitored and kept at 37–38 °C using a closed-loop heating pad, and the eyes were protected with ophthalmic gel (Viscotears Liquid Gel, Alcon). An analgesic (Rimadyl, 5 mg/kg) was administered subcutaneously before the procedure, and orally on subsequent days. Dexamethasone (0.5 mg/kg, IM) was administered intramuscularly 30 min before the procedure to prevent brain oedema. The exposed brain was constantly perfused with artificial cerebrospinal fluid (150 mM NaCl, 2.5 mM KCl, 10 mM HEPES, 2 mM CaCl2, 1 mM MgCl2; pH 7.3 adjusted with NaOH, 300 mOsm). During the surgery, we first implanted a head-plate over the right hemisphere of the cranium for later head-fixation: a stainless-steel head plate with a 10-mm circular opening was secured over the skull using dental cement (Super-Bond C&B, 10 Sun Medical). We then made a circular craniotomy over V1 (3 mm diameter) using a biopsy punch. At this point 6-7 virus injections were made at different positions inside the craniotomy. Finally, the craniotomy was sealed with a glass cranial window, using cyanoacrylate adhesive (Vetbond, 3M) and dental cement.

All mice were injected with an unconditional GCaMP6m virus, AAV1.Syn.GCaMP6m.WPRE.SV40 obtained from the University of Pennsylvania Viral Vector Core. The virus was injected with a bevelled micropipette using a Nanoject II injector (Drummond Scientific Company, Broomall, PA 1) attached to a stereotaxic micromanipulator. Six to seven boli of 100-200 nL virus (2.23×10^12^ GC/ml) were slowly (~20 nL/min) injected unilaterally into monocular V1^82^, 2.1-3.3 mm laterally and 3.5-4.0mm posteriorly from Bregma and at a depth of L2/3 (200-300 mm).

After virus injection, a small bolus (10uL) of red fluorescent beads (FluoSpheres™ Carboxylate-Modified Microspheres, 2.0 μm, red fluorescent (580/605), 2% solids, ThermoFisher Scientific) was injected at the most rostral part of the craniotomy, to allow orientation of the ex-vivo slices but not interfere with V1 imaging in the caudal part. Following recovery, mice were habituated for handling and head-fixation for 3 days before carrying out recordings.

### Recording neuronal activity in V1

#### Two-photon calcium imaging

Each mouse was recorded for at least 3 sessions. *In vivo* recordings were performed 15-45 days after the virus injection. We used a commercial two-photon microscope with a resonant-galvo scanhead (B-scope, ThorLabs, Ely UK) controlled by ScanImage 4.2^83^, with an acquisition frame rate of about 30 Hz (at 512 by 512 pixels, corresponding to a rate of about 4.3 Hz sampling rate). The field of view was 550-600 μm large. We imaged 7 planes at 15-45 μm steps, starting at various positions below the brain surface (from 0 to −150 μm) to sample different cortical depths and therefore t-types recorded simultaneously during different sessions. Imaging calcium activity was performed at a wavelength of 920nm or 980nm. Three computer screens spanning −135 to +135 v° along the azimuth axis and −35 to +35 v° along the elevation axis were used to display visual stimuli. During the presentation of visual stimuli, we switched off the red gun of the monitors to prevent light from the monitors contaminating the red fluorescent channel.

At the end of each recording session, reference Z-Stacks were acquired. Starting at the same position as the imaging planes, we acquired two Z-Stacks of about 400um depth, with a 1-micron step between planes. The first one, called GCaMP Z-Stack was acquired at the same wavelength as the calcium imaging (920 or 980nm). The second one, called reference Z-Stack, was acquired at 1040nm to image mCherry fluorescence.

Before sacrificing each mouse, we acquired structural Z-stacks (ranging from the brain surface to 400um deep) at 1040nm to get an image of the mCherry cells across the whole craniotomy (including the position where the red fluorescent beads were injected). This structural Z-stack was used to select slices on which to perform transcriptomic analysis, and to provide an initialization point for the registration algorithm.

#### Initial retinotopic mapping

All recordings were targeted to the V1 Monocular region (>60° azimuth). To find this region, during the first imaging session, we initially mapped the retinotopy of different candidate fields-of-view, using single plane imaging. Sparse noise stimuli were presented to the mouse, consisting in black or white squares of width 4.5° visual angle on a grey background at a frame rate of 5 Hz for 10 minutes. Squares appeared randomly at fixed positions in a 16 by 60 grid, spanning the retinotopic range of the computer screens. 1.5% of the squares were shown at any one time.

#### Visual Stimulation

Drifting gratings were centred on the mean receptive field of the microscope’s field of view. Gratings had a duration of 0.5 s, temporal frequency of 2 Hz and spatial frequency of 0.15 cycles/deg. The gratings drifted in 12 different directions (from 0 to 330°, separated by 30°) and were of 3 different sizes (5°, 15° and 60° diameter).

Natural scenes from the ImageNet database were contrast-normalized and presented as described in Ref.^49^. Each image was presented for 0.5 s with inter-stimulus interval uniformly distributed from 0.3 to 1.1 s. Five percent of the total presentations were blank stimuli. During each session we presented a given set of 1050 different natural images twice (corresponding to a subset of the 2800 images originally used in Ref.^49^).

On each recording session we presented the same random sparse noise stimuli used to map retinotopy (see above), for 30 minutes.

Spontaneous activity was recorded in front of a blank screen, set to a steady cyan level equal to the background of all the stimuli presented for visual responses protocols. The duration of these blank screen presentations was typically between 15 and 20 minutes long.

### Eye-Tracking

We used a collimated infrared LED (SLS-0208-B, lpeak = 850nm; controller: SLC-AA02-US; Mightex Systems, Toronto, Canada) to illuminate the eye contralateral to the recording site. Videos of eye position were captured at 30 Hz with a monochromatic camera (DMK 21BU04.H, The Imaging Source, Bremen, Germany) equipped with a zoom lens (MVL7000; Navitar, Rochester, NY), and positioned at approximately 50 degrees azimuth and 50 degrees elevation relative to the centre of the mouse’ field of view. Contamination light from the monitors and the imaging laser was rejected using an optical band-pass filter (700-900nm) positioned in front of the camera objective (long-pass 092/52×0.75, The Imaging Source, Bremen, Germany; short-pass FES0900, Thorlabs, Ely UK).

### Processing of calcium imaging

Two photon calcium data was processed using Suite2P^84^. Neuropil contamination was corrected by subtracting from each ROI signal its surrounding neuropil signal multiplied by a constant factor of 0.7. Calcium traces were deconvolved using non-negative spike deconvolution^85^ with a calcium indicator decay timescale of 1.5 s. ROIs were manually curated to make sure only cell bodies were considered for further analysis.

### coppaFISH: Combinatorial Padlock-Probe-Amplified Fluorescence *in Situ* Hybridization

Many approaches to highly-multiplexed mRNA detection have been described^42,43,86–102^. The coppaFISH method is a development of the *in situ* sequencing method of Ref.^41^ (Extended Data Fig. 1). The method uses reverse transcription, padlock probes, and rolling-circle amplification to amplify mRNAs to DNA rolling circle products (RCPs) containing multiple copies of a 20 nucleotide (nt) barcode sequence, and then detects their location combinatorially in 7 rounds of 7-colour fluorescence imaging.

#### Gene selection and DNA probe design

A panel of 73 genes was selected to allow the identification of cortical cell types. This gene panel was essentially the same as the one used in Ref.^41^., except that some genes that were previously found not to help cell type identification were removed. One gene (*Yjefn3*) was detected in our experiments, but could not be used to assign cells to transcriptomic t-types, as it was not present in the reference scRNA-seq dataset^4^. In the main text we therefore refer to a 72-gene panel.

Multiple padlock probes were designed for each gene, spanning the length of the cDNA (Supplementary Data File 2). The number of different padlock probes per gene was chosen based on the expression for each specific gene as determined by scRNA-seq. This means that fewer padlock probes were used for genes with low expression and vice versa (for example 4 padlock probes were designed for Sst but 10 were designed for Chodl). All padlock probes consisted of two 15-20nt recognition sites, a 20nt gene barcode (unique to each gene) and a 20nt anchor sequence (identical for all genes and padlock probes).

Padlock probes were designed using the software of Ref.^41^. Briefly, this software finds suitable RNA target sequences by restricting the melting temperature of the binding sequence, and by aligning the candidate sequences to the mouse whole transcriptome (RefSeq database) using BLAST+ to check for specificity. Any candidate targets for which another transcript or non-coding RNA sequence matched the target with more than 50% coverage, 80% homology, and coverage spanning the central 10nt of the target sequence were excluded. For each padlock probe we also designed a specific primer for reverse transcription, a 15nt long DNA oligonucleotides which binds the region upstream to the mRNA sequences targeted by the padlock probes (Supplementary Data File 3). The use of specific primers greatly improved the number of RCPs obtained per section compared to random primers (our unpublished observations).

To determine the gene-specific DNA barcode sequences (and the anchor sequence), 240,000 orthogonal 25-mer oligonucleotide sequences^103^ were trimmed to 20nt from the 5’ end and screened for Tm (between 55 and 56 °C using SantaLucia method). They were further screened for orthogonality with mouse sequences using BLAST+ with the NCBI mouse genomic plus transcript (Mouse G +T) database. We used the following BLAST parameters: “-reward”, 1, “-penalty”, −2, “-gapopen”, 2, “-gapextend”, 1, “-evalue”, 10. Any matches in this blast search were removed from the pool. Next, we checked for potential cross reactivity of the remaining sequences to themselves using the same BLAST parameters, and any hits were removed, resulting in 6397 possible sequences. The barcode sequences were chosen from this pool.

The combinatorial imaging strategy used two types of DNA Probes. Seven “Dye probes” were designed, each consisting of a 20nt long DNA oligo conjugated to one of the 7 following fluorophores: DY405, AF488, DY485xL, AF532, AF594, AF647 and AF750; the same dye probes were used on each imaging round (Supplementary Data File 4). Additionally, a set of 40nt “Bridge probes” were designed for each imaging round, linking each gene’s RCP barcode to one of the 7 Dye probes (Extended Data Fig. 1; Supplementary Data File 5). These bridge probes thus caused each gene to show up in a specific colour channel on each round. This two-part strategy of linking the 7 dye probes to the RCPs with bridge probes provides a substantial cost saving over making *N_genes_* × *N_rounds_* dye probes, as dye-coupled probes are much more expensive than simple DNA.

Each gene was assigned a sequence of dyes for the 7 imaging rounds using a Reed-Solomon coding scheme^104^ (Supplementary Data File 6), which constructs sequences of minimum possible overlap. Specifically, the genes were numbered by integers *g,* and converted to a base 7 representation *g*_2_*g*_1_*g*_0_. The dye assigned to gene *g* on round *r* was

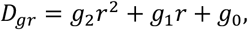

where addition and multiplication are understood to happen modulo 7. Codes 0 to 6, which correspond to the same colour in each round, were not used as these codes could not be distinguished from fixed background fluorescence.

All custom DNA oligos (Padlock probes, primers, Bridge probes and Dye probes) were obtained from Integrated DNA Technologies (Leuven, Belgium). Padlock probes were ordered as 5’ phosphorylated 4 nmole Ultramer™ DNA oligos, all other oligos were ordered as classical 25 nmole DNA oligos. The DNA sequence for all 556 primers and padlock probes, 511 bridge probes and 7 dye probes are provided in (Supplementary Data File 2-5).

#### Tissue preparation

After the *in vivo* recordings were finished, mice were anaesthetized with isoflurane and then injected with a lethal dose of sodium pentobarbital (0.01 ml/g). The fresh brains were then dissected out from the skull taking great care to preserve the integrity of the tissue and avoid warping. The brains were then placed in OCT (Sakura Finetek) and left to freeze on dry ice for 30min. The samples were then stored at −80°C until slicing. 15-μm thick sagittal sections were then obtained using a Leica Cryostat for each brain and mounted on gelatine-coated borosilicate glass coverslips (22×55mm). Gelatine coated coverslips allowed tissue section adhesion to the coverslip and RNA preservation throughout the protocol. To make them, coverslips mounted on a rack were dipped for 30 seconds in solution of a 2% w/v gelatine and 0.2 % w/v chromium potassium sulphate dodecahydrate in distilled water (https://www.rndsystems.com/resources/protocols/protocol-preparation-gelatine-coated-slides-histological-tissue-sections). 2-3 brain sections were thaw-mounted on each coverslip and then frozen and stored at −80°C.

#### In situ rolling circle product (RCP) production

The RCPs were prepared as in Ref.^41^, with some modifications. First, coverslips were taken out of the freezer and then directly pre-fixed using 4% PFA for 5 minutes at room temperature. This pre-fixation was followed by a quick wash with nuclease-free PBS, and incubation in 0.1 M HCl for 5 minutes at room temperature. After one more PBS wash, the sections were incubated in 70% Ethanol for 1 minute and then in 100% Ethanol for 1 minute at room temperature. The coverslips were then left to dry in air. To keep the reagents on the tissue sections, a barrier was drawn around each section using a hydrophobic barrier PAP pen (ImmEdge® Hydrophobic Barrier PAP Pen H-4000 - Vector Laboratories).

The sections were then directly incubated in reverse transcription mix overnight at 37°C in a humidified chamber (Slide staining system, StainTray™ M918, VWR^TM^). The mix contained 0.5 mM dNTP mix (Thermo), gene specific primers (10 μM each), 0.2 μg/μL BSA (NEB), 1 U/μL RIBOPROTECT RNase Inhibitor (Blirt) and 20 U/μL TranscriptMe reverse transcriptase (Blirt) in 1x reverse transcription buffer (Blirt). The mix was removed and fresh 4% (w/v) paraformaldehyde in PBS was added to the sections without any wash in between. This post-fixation step aimed to cross-link newly synthesized cDNA to the cellular matrix and was carried out at room temperature for 30 minutes, followed by two washes in PBS. RNaseH digestion, padlock hybridization and ligation were then performed using a single reaction mix. The mix contained 0.05 M KCl (Sigma), 20% Ethylene Carbonate (EC) (Sigma), 10 nM of each padlock probe (557 probes), 0.2 μg/μL BSA, 0.3 U/μL Tth DNA Ligase (Blirt) and 0.4 U/μL RNase H (Blirt) in 1x Ampligase buffer (epicenter). The sections were first incubated at 37°C for 30 min for RNaseH digestion and moved to 45°C for 60 minutes for stringent hybridization and optimal DNA ligase activity. The sections were then washed twice in PBS. Finally, for rolling circle amplification, the sections were incubated in a mix containing 5% glycerol (Sigma), 0.25 nM dNTP mix, 0.2 μg/μL BSA, 0.2 U/μL EquiPhi29 DNA Polymerase (Thermo Fisher Scientific) and 1x EquiPhi29 buffer (Thermo Fisher) overnight at 30°C.

RCP production was quickly verified prior to full barcode read-out by hybridizing a AF750-conjugated oligonucleotide probe (IDT) to the anchor sequence present in all the RCPs. Sections were incubated for 15 minutes at room temperature in a hybridization mix containing 10 nM of the dye probe, 2xSSC, 20% EC and H2O. They were then washed twice with 2xSSC. The SSC was then removed from the sections and the coverslips were mounted onto SuperFrost plus (VWR) glass slides using 10 μL SuperFrost gold antifade mountant (Life Technologies). Images of the region of interest (visual cortex) were then acquired to visualize the RCPs.

#### Imaging of the in situ barcodes (read-out)

All seven rounds of imaging occurred in a custom flow cell, using automated fluidics to wash appropriate bridge and dye probes prior to each round. The flow cell frame was designed using *Blender* and printed, using an Ultimaker S5 3D printer, in polylactic acid filament (PLA) with polyvinyl alcohol (PVA) support structures. The PVA support was removed after printing by placing the flow cells in water on a rocker overnight. To make the flow cell air-tight, two 22×55 mm glass coverslips (one with RCP containing sections and one bare) and two approx. 40 cm long EFTE tubes (Tubing Tefzel Nat 1/16 OD x .020 ID) were securely mounted using UV curing cement (Norland Optical Adhesive 81) and UV curing LED system with driver unit and handheld 365nm light source (ThorLabs, CS20K2). The coverslip with the sections was mounted so that the side with the sections faces the inside of the flow cell.

The Imaging setup consisted of a Nikon Eclipse Ti2 microscope with a NIR-LDI laser panel and a zyla sCMOS 4.2 camera (Andor). The fluidics setup consisted of a Minipuls 3 pump (Gilson) and two linked MVP multivalves (Hamilton), each with 8 ports. Nikon NIS elements software was used to acquire the images and communicate with a second computer controlling the fluidic pump and multivalves. The opening of the valves and the speed and the duration of the pump’s activity was managed by an edited version of Kilroy software (https://github.com/ZhuangLab/storm-control; edits available at https://github.com/acycliq/storm-control). The imaging and sequencing chemistry were coordinated by NIS elements software (ND sequence acquisition module), which communicates with the computer running Kilroy by sending TTL pulses through a National Instruments NI-USB 6008 board.

Before sequencing, 15 mL falcon tubes containing bridge probe mixtures for each of the seven imaging rounds, as well as one each for dye probe mixture, anchor probe mixture, imaging buffer, distilled water, 2xSSC, and 100% formamide were attached to the multivalves via EFTE tubing and flangeless fittings (1/16 inch Red Delrin, IDEx Health and Science LLC). The mixtures for bridge, dye, and anchor probes contained the appropriate oligonucleotides diluted to 10nM each in 2xSSC, 20%EC, and H2O. The bridge probe mix for the final anchor round contained the Cy7-conjugated anchor probe as well as the Gad1 bridge probe (10nM) that binds to the AF532 dye probe (Gad1_r6 - 10nM) and DAPI to stain the cell nuclei. A fresh formamide (S4117 Millipore) aliquot was used for every experiment (stored at 4 °C). The flow cell was then mounted onto the multi-slide stage and connected to the pump and multivalves via EFTE tubing. The speed of the pump was adjusted to approximately 0.4 mL/sec. To fill the flow cell, each solution was flushed through the fluidics system for 4 minutes (the flow cell volume is approximately 1 mL).

In total, eight rounds of imaging were done for each imaging experiment: 7 rounds to decode the barcodes and one final anchor round to detect the position of every RCP that was used for later image alignment. In each round, sections were first incubated in 100% formamide for 15min to strip the RCPs from any previous labelling. The formamide was then flushed from the flow cell with water for 4 minutes and then with 2xSSC for 4 minutes. The sections were next incubated in that round’s bridge probe mix for 15 minutes and washed with 2xSSC. After this, the sections were incubated in the dye probe mix for 15 minutes, and again washed with 2xSSC. The flow cell was filled up with an imaging buffer consisting of glucose oxidase and catalase containing oxygen scavenging system^105^ to protect the fluorophores from photobleaching during imaging.

After each round of sequencing chemistry, 16-bit images were acquired using wide-field epifluorescence excitation, and a 40X magnification air-objective (CFI Plan Apochromat Lambda 40XC - NA 0.95). Images consisted of Z-stacks (z-step: 0.5um) in 7 different colour channels corresponding to the 7 fluorophores (Fluorophore – excitation wavelength, emission filters: Dy405 - ex405, 460/50m; AF488 - ex470, 525/36m; Dy485xl - ex470, 632/60m; AF532 - ex520, 560/40m; AF594 - ex555, 632/60m; AF647 - ex640, 700/75m; AF750 - ex730, 811/80m). Each tile was 2048×2048 pixels (pixel size: 0.1625 micron). The imaging parameters were adjusted to cover only the region of interest (V1) and usually consisted of 10-15 tiles with 10% overlap. The Nikon perfect focus system was used to make sure that the focus stayed relatively constant across imaging rounds. Image files were saved in Nikon’s native ND2 format.

### *In situ* data analysis

The *in situ* data was analysed with a suite of custom software for image processing, gene calling, and cell calling. All code was written in MATLAB, and is freely available at https://github.com/jduffield65/iss. This software was developed from that described in Ref.^41^, but has been greatly modified, so is described in full here.

The *in situ* data consist of 8 rounds of multispectral imaging (7 combinatorial rounds, and one reference round in which all RCPs are labelled via the anchor sequence, together with an additional stain for Gad1 RCPs and a DAPI stain). Because the tissue sample is too large for a single camera image, imaging occurs in overlapping tiles. In each tile, a focus stack of widefield images were taken for each colour, and flattened into 2D using an extended depth of focus algorithm^106^. The data therefore consists of a set of images

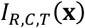

Here *I* gives the pixel intensity for sequencing round *R,* colour channel *C,* tile *T,* and pixel coordinates x within this tile. The processing pipeline to identify detected genes comprises several steps: initial registration; RCP spot detection and fine registration; crosstalk compensation; and gene calling. These analyses proceed without ever “stitching” all the tiles into a single large image; this approach allows processing of very large datasets on computers with limited memory, and also easily allows non-rigid alignments. Prior to the pipeline, all RCP images are linearly filtered by convolving with a difference of Hannings: a Hanning of radius 0.5 μm minus a Hanning of radius of 1 μm, both normalized to have sum 1. The DAPI background images are filtered with a disk-shaped top-hat filter with radius of 8 μm.

#### Initial registration

The initial registration step finds offsets between all image tiles using the anchor images taken on round 8 (which we refer to as “reference images”). We use this to define a global coordinate system for the entire tissue sample.

Because we use a square tiling strategy, each tile may have up to four “neighbours”: other tiles with which it has a region of substantial overlap. We denote the set of neighbouring tile pairs as 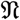.

Spots first are detected in each tile’s reference images, as local maxima of the filtered image exceeding a fixed detection threshold. To align the reference images, we loop over all pairs of neighbouring tiles, and compute an offset to register the overlapping regions of the filtered reference images of these two tiles. The offset between two tiles *T*_1_ and *T*_2_ is found by exhaustive search over all 2d shifts in a range around to the shift expected from the microscope’s position sensor. For each shift, we find for each spot *s* on *T*_1_ the pixel distance *D_s_* to the nearest spot on *T*_2_ after the shift has been applied. A score is computed as ∑_*s*_ *e*^−*D*_*s*_2/8^, and the final shift vector Δ_*T*_1_,*T*_2__ is taken as the one maximizing this score i.e. the one with the most near neighbours.

We define a single global coordinate system by finding the coordinate origin **X**_*T*_ for each tile *T*. Note however that this problem is overdetermined as there are more neighbour pairs than there are tiles. We therefore compute the offsets by minimizing the loss function

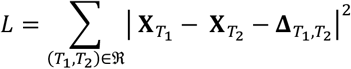

Differentiating this loss function with respect to **X**_*T*_ yields a set of simultaneous linear equations, whose solution yields the origins of each tile on the reference round. The results of this step suffice to define a global coordinate system, but do not provide pixel-level alignment of images from multiple colour channels on multiple rounds, due to the occurrence of chromatic aberration and small rotational or non-rigid shifts. The latter will be dealt with in the next step, through point-cloud registration.

#### Spot detection and fine registration

The second processing step detects spots in all images of the 7 sequencing rounds, performs fine alignment of colour channels and sequencing rounds, and computes for each spot a position in global coordinates and an intensity vector summarizing that spot’s detected fluorescence in each round and channel.

The most intricate part of this step is fine image registration. Even though the same tile layout is used for all sequencing rounds, the precise positions of the tiles may differ due to slight shifts in the placement and rotation of the sample. Thus, a single spot might be found on different tiles in different sequencing rounds. Furthermore, due to chromatic aberration a spot may be in slightly different positions (although not different tiles) in different colour channels. Because most spots are only a few pixels in size, even a one-pixel registration error can compromise accurate RNA reads.

A global coordinate is defined for each of the spots detected in the reference images using the initial registration described above. In regions where tiles overlap, duplicate spots are rejected by keeping only spots which are closer in global coordinates to the centre of their original tile than to any other.

Next, spot positions are detected in images from all sequencing rounds and colour channels. These are used to align each round and colour channel to the corresponding tile’s reference image, using point-cloud registration. Specifically, we fit an affine transformation from each reference image to the images of the corresponding tile for all rounds and colour channels, using the iterative-closest point (ICP) algorithm with matches further than 3 pixels away excluded. These affine transformations can include shifts, scalings, rotations and shears, but we did not find it necessary to introduce nonlinear warping transformations within tiles (nonlinear transformations can still occur globally by variation of the affine transformation across tiles). As the ICP algorithm is highly sensitive to local maxima, it is initialized from a shift transformation computed by the same method used to find the overlap between reference images, i.e. the shift that maximizes the number of near neighbours as measured by ∑_*s*_ *e*^−*D*_*s*_2/8^. When spots are located on neighbouring tiles on different rounds, the corresponding images are again registered with ICP.

Finally, a 7-dimensional intensity vector **v**_*s,r*_ is computed for each spot s in each round *r,* by reading the intensity from the aligned coordinate of each filtered image.

#### Crosstalk compensation

The last step associating spots to genes consists of transforming the intensity vectors to gene identities.

An important consideration in this stage is that crosstalk can occur between colour channels. Some crosstalk may occur due to optical bleedthrough; additional crosstalk can occur due to chemical cross-reactivity of probes. With the current hybridization chemistry (unlike previous sequencing-by-ligation chemistry), the degree of crosstalk tends to be constant within a round, so we learn a single 7×7 crosstalk matrix and apply it to all rounds.

To estimate the crosstalk present, we first collect a set of seven 7-dimensional vectors **v**_*s,r*_ containing the intensity in each colour channel of all well-isolated spots s in all rounds *r.* Only well-isolated spots are used to ensure that crosstalk estimation is not affected by spatial overlap of spots corresponding to different genes; a spot is defined as well-isolated if the reference image intensity averaged over an annular region (4-14 pixel radius) around the spot is less than a threshold value. Crosstalk is then estimated by running a scaled k-means algorithm ^107^ on these vectors, which finds a set of seven vectors ***c**_d_* (*d* refers to one of the seven dyes), such that the error function:

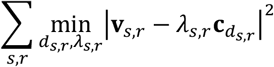

is minimized; in other words, it finds the seven intensity vectors **x**_*c*_ such that each well isolated spot on round *r* is close to a scaled version of one of them.

The crosstalk matrix is used to predict the colour profile expected for an RCP of each gene *g,* for each colour channel and round. If gene *g* is assigned the dye *d_gr_* in round *r,* the predicted 49-dimensional intensity vector is obtained by concatenating the corresponding crosstalk vectors

#### Gene calling

Improvements in tissue processing and *in situ* chemistry mean that our current methods produce substantially more RCPs than the previous *in situ* sequencing method^41^. Consequently, the fluorescence of neighbouring RCPs often overlaps, which would render the previous detection method unable to find them. To allow resolution of overlapping spots, we therefore developed a new gene calling algorithm, based on orthogonal matching pursuit (OMP)^107^. This algorithm also allows for subtraction of background autofluorescence. Essentially, OMP repeatedly tests whether the 49-dimensional fluorescence vector of a pixel overlaps with the predicted fluorescence vector of each gene; if so, a gene is detected at that location, its code is projected out from the fluorescence vector, and the process repeats.

The OMP algorithm fits a 49-dimensional image (one dimension for each combination of round and colour channel) as a sum of 49-dimensional code vectors. There is one code vector **a**_*g*_ for each gene, and one “background” code 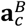 for each colour channel, which has equal intensity for all rounds in one colour channel only. These background codes account for tissue autofluorescence, which will affect all imaging rounds equally.

The gene codes **a**_*g*_ are derived from the using knowledge of the Reed-Solomon assigned dyes *d_g,r_* for each gene in each round and the crosstalk matrix columns **c**_*d*_. These codes take into account the fact that different genes can have consistently different intensities in different rounds, which may arise from non-uniformity in the synthesized concentrations of the bridge probes. To account for this non-uniformity, we learn a scale factor *ε_g,r_*, and predict the 49-dimensional gene code for gene *g* as a concatenation:

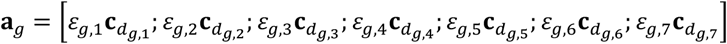

We will describe the general algorithm before specifying how *ε_g,r_* is chosen.

The OMP algorithm expresses the 49-dimensional fluorescence vectors **v**_*p*_ for each pixel *p* as a weighted sum of code vectors: 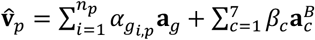. Each step of the algorithm can add a code to the set of code vectors {*g_i,p_*: *i* = 1 … *n_p_*} used to approximate pixel **v**_*p*_; the 7 background codes are always included. The gene set is initialized to be empty, and to choose which gene code, if any, should be added on each step the algorithm computes how much the residual 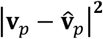 would decrease for each possible addition to the set, and picks the gene giving maximum decrease, provided this decrease is above a threshold of 0.0612 multiplied by the second largest absolute value of **v**_*p*_ (clamped by a minimum threshold of 0.01 and a maximum threshold of 3.0), up to a total of 6 genes per pixel. After this iterative process has terminated for all pixels, an image is made for each gene, containing the gene’s weight for each pixel or zero if that gene is not in the pixel’s gene set. RNA detections are found as local maxima of this image, subject to a thresholding criterion; the criterion takes into account several factors and is best understood by examining the source code (https://github.com/jduffield65/iss).

To choose the scale factors *ε_g,r_*, a single iteration of the OMP algorithm is run with all *ε_g,r_* = 1. Local maxima detected as just described, but with a more stringent threshold (see source code for details) to ensure only unambiguous gene detections are used. We then compute a 7-dimensional mean intensity vector 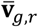 of all detected spots for each gene in each round. We then find the scale factors *ε_g,r_* for each round and gene as the least-squares solutions of

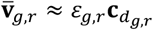

#### Cell calling

The DAPI image was used to segment the cells. This was performed by detection of the local maxima in each cell followed by watershed segmentation. The segmentation of matched cells and their close neighbours was manually curated.

To assign cells to transcriptomic t-types, we used the pciSeq algorithm of Ref.^41^, a Bayesian method which assigns each *in situ* cell a posterior probability of belonging to each of a set of cell classes defined by prior scRNA-seq. The cortical t-types and their mean gene expression were obtained from Ref.^4^ using only V1 data. The read counts of this scRNA-seq data were divided by 100 to predict the expected *in situ* RNA count; a further gene-dependent efficiency factor was estimated by the algorithm. The pciSeq algorithm produces a probability for each cell to belong to each class, which we converted to a “hard” classification by assigning each cell to the t-type of maximum *a posteriori* probability; cells for which this maximal probability was less than 0.5 were not analysed further (~1% of matched cells).

### Registration of the in vivo and ex vivo cells

We used inhibitory cells, labelled *in vivo* by mCherry (Gad2-mCherry mice), as landmarks to perform the registration between the *in vivo* Gad-mCherry volume and the *ex vivo* brain sections (Extended Data Fig. 2). This alignment made use of two high-resolution reference Z-stacks taken for each subject following each imaging session. The “GCaMP Z-stack” was taken using the same wavelength as functional imaging (920 or 980nm), covering the same volume but at higher resolution. The “mCherry Z-stack” was acquired in the same volume with 1040nm excitation wavelength to detect inhibitory neurons in Gad2-mCherry mice, but also provided some GCaMP signal in the green channel (although this signal was much lower than for the GCaMP Z-stack taken at 920nm). The different excitation wavelength of these two Z-stacks led to a small chromatic aberration, which was only significant in depth. To correct this aberration, we used the green channel found in both imaged volumes, registering planes of the GCaMP Z-stack to the mCherry Z-stack using FFT convolution. This was achieved by finding the best matching plane from the later Z-stack for each GCaMP Z-stack planes as the Z position which gave the highest FFT cross-correlation. Additionally, a “global Z-stack” was made following the final functional imaging session, covering the entire region under the craniotomy, used for coarse initial registration of the *in situ* slices.

#### Aligning Calcium ROIs to the mCherry Z-Stack

To align the imaging planes of one functional 2p session to the GCaMP Z-stack, we first obtained their theoretical position using the measured position of the objective for each line scanned (for both the functional imaging planes and the GCaMP Z-Stack). We then estimated the Z-drift during the recording session: the position of the calcium imaging planes over time in comparison to this GCaMP Z-Stack. To do so, a mean image of each functional imaging plane was obtained for 1 minute every 7 minutes of the recording. These mean images were then aligned to the Z-stack using FFT (Fast Fourier Transform) convolution. We then took the median of this Z-drift over time and used it to correct the theoretical imaging plane position. We then performed FFT based registration to correct for a small shift in X and Y between the actual mean image and the reconstructed image. We thus found the position of the imaging planes (and therefore of each functional ROI) in the GCaMP Z-Stack. These were then aligned to the mCherry Z-Stack using the transformation described above (chromatic aberration in depth).

#### Aligning brain slices to the mCherry Z-Stack

To register the positions of the *in situ* detected inhibitory neurons to the 3D mCherry Z-stack, we used a custom point cloud registration method, using inhibitory neurons as landmark points. MATLAB code and an example pipeline script can be found at https://github.com/ha-ha-ha-han/NeuromicsCellDetection/.

During slicing, the latero-medial order of the sagittal brain sections was carefully recorded. To find the sections corresponding to the imaged region, we first screened them by generating RCPs for every 20th section, and staining with the Gad1 bridge probe and its corresponding dye probe to label inhibitory neurons. The position of the fluorescent bead injection was usually visible on one of the sections, allowing us to infer the approximate position of every slice (based on the known order and thickness of slicing).

Fine registration of screened sections to the *in vivo* reference Z-stack started with cell detection *in vivo and ex vivo*. To detect cells in the *in vivo* mCherry Z-stack, each plane was contrast normalized to correct for the loss of brightness with depth using the following MATLAB GUI https://github.com/nadavyayon/Intensify3D/blob/master/User_GUI_Intensify3D.m, which performs background and signal estimation based on user defined thresholds), and the Z-stack was then filtered using a 3D median filter of radius 2 μm to reduce background noise. The mCherry positive cells were automatically detected on these images using a 3D difference-of-Gaussians filter followed by watershed segmentation. Manual curation was performed to correct for missed or false positive detections. To detect inhibitory cells in the *ex vivo* slices, we used the *in situ* expression of Gad1 in the reference round, since native mCherry fluorescence was not preserved in the fresh-frozen sections. Gad1 detections formed clusters on GABAergic cells (Extended Data Fig. 2), which were detected by Gaussian smoothing of the Gad1 RCP images and applying a difference of gaussian filters and watershed segmentation to detect individual clusters. Finally, we manually curated these detections using the full *in situ* 72 gene expression to determine putative interneurons based on the main inhibitory cell markers such as Vip, Sst, Pvalb etc.

The slices were first coarsely registered using brain structures (hippocampus, brain surface etc.) visualized using the anchor and nuclear staining. Next, they were finely registered using an algorithm to register a 2D point cloud corresponding to inhibitory neurons in the *ex vivo* slice into a 3D point cloud corresponding to inhibitory neurons in the *in vivo* volume. To align these clouds, we used rigid registration with 6 degrees of freedom (*α, β, γ, x, y, z*), where *α, β, γ* are the rotation angles, and *x, y, z* are translational shifts. (Non-rigid point cloud registration is possible, but we found it to be unnecessary.) The registration algorithm searched for the parameters (*α_max_, β_max_, γ_max_, x_max_, y_max_, z_max_*) that maximize the match of the 2D slice to the corresponding section of the 3D volume.

Because this registration problem has a large number of local maxima, we performed an exhaustive grid-search over these 6 parameters. Because Fourier convolution of 3d arrays is fast, but rotation of them is not, we used a hybrid point/Fourier method. An outer loop searches over all combinations of rotation angles (*α, β, γ*), with an initial step size of 1°, refined to 0.5° for finer alignment, and rotates the 3d point cloud accordingly. A 3d volumetric image is then synthesized from these rotated points by adding a Gaussian peak at the location of each point. Each plane *z* of this image is Fourier convolved with a fixed 2d array synthesized similarly for the 2d cloud, and the resulting 3d correlation map is stored, to accumulate a correlation score function *c*(*α, β, γ, x, y, z*). The top local maxima of this 6d array are found and ranked using both the intensity of the cross-correlogram peaks and the percentage of cells matched within a tolerance of 15 microns (to account for small non-rigid deformations). Finally, the match validity for each section was assessed manually by looking at the overlay between the interpolated cut from the reference Z-Stack and the Gad1 RCP image. The rotation and translation parameters were manually adjusted to provide the best overlay between the two datasets. Typical rotation angles were found between −10 and 10 degrees of the coarse manual registration, enabling us to save computation time by searching only this range.

#### Aligning individual neurons

Finally, a custom MATLAB GUI was used to curate the match between inhibitory cells in the *in vivo* recordings and the *ex vivo* sections. The GUI allowed us to visualize the *in vivo* mCherry image of each cell (obtained from the reference Z-Stack), the position of the ROIs on the reference Z-Stack and the overlap between the reference Z-Stack cross-sections and the *in situ* gene expression for the different genes. For each slice, we displayed all mCherry positive ROIs which were less than 20 μm away from the found position of the slice in the reference Z-Stack. Each assignment of *in vivo* and ex-vivo Gad positive cells was curated manually based on this data. At this stage the boundaries initially found using automatic segmentation of the DAPI image were also manually adjusted for the matched cells and their neighbours, to correct for errors in DAPI segmentation that could impact the gene and cell type assignment. This correction was based both on the DAPI image and on the *in situ* gene expression, which provided information that could indicate under-splitting in the DAPI segmentation of adjacent cells.

### Class and Cell selection

We recorded a total of 3469 (204±42 per session) inhibitory cells and together with 6684 (393±173 per session) excitatory cells. Of these inhibitory cells, we managed to match with good confidence and assign a t-type to 1515 (89±30 per session) cells (see Supplementary Data File 1). Some *ex-vivo* identified cells were recorded in multiple imaging sessions. In all figures a unique session was picked for each matched cell (except Fig. 2 where we show all cells in a single session). The session assigned was chosen based on the percentage of time the mouse spent running during this session, to maximize variability of behaviour while the cell was recorded. After removing these duplicates, we obtained 1028 unique cells. Finally, 8 cells which were assigned to t-types with less than 3 cells total were discarded. The final population of 1020 cells belonged to 35 transcriptomic t-types.

For hierarchical analysis, the 35 t-types were grouped into 11 Classes corresponding to putative anatomical/physiological cell types based on the previous literature. For Pvalb neurons, the grouping was unambiguous: the Pvalb-Vipr2 t-type is genetically very different to all other Pvalb t-types, and several studies have identified molecular markers of this t-type with chandelier cells^4,6,8,108^. For Sst cells, UMAP analysis (Extended Data Fig. 3) suggests that the two Sst-Tac1 t-types bridge a continuum between the two Sst-Calb2 t-types (identified as superficial-layer Martinotti cells^8,109,110^) and the Pvalb-Tpbg t-type (identified as superficial-layer Pvalb basket cells^8^). Patch-seq analysis confirms that Sst-Tac1 cells have less axon in L1 and faster-spiking phenotypes than classical Martinotti cells^8^. We therefore identify the two Sst-Tac1 t-types as non-Martinotti Sst cells, acknowledging that these two Sst Classes likely tile a continuum, rather than truly being discrete cell types. For Lamp5 cells, we grouped t-types based on the results of Ref.^111^ (see also Ref.^112^). The three t-types comprising the Lamp5-Npy group were identified as neurogliaform cells based on their strong expression of Npy. The Lamp5-Fam19a1-Tmem182 t-type was identified as Canopy cells due to expression of Ndnf but not Npy; the two remaining t-types were identified as *α*7 cells due to their strong expression of Chrna7 and weak expression of Ndnf and Npy. For Vip cells, we divided t-types by transcriptomic methods: UMAP analysis suggested a clear discrete distinction between two Vip t-types characterized by expression of Reln as well as weaker expression of Vip itself. We are not aware of any specific study on these Vip-Reln cells, however based on their weak Vip expression, the fact that Reln is a usually L1 marker, we provisionally identify this Class with the layer 1 VIP cells described by Ref.^111^. Serpinf1 t-types were included with the Vip category as we do not see strong evidence for this as a discrete Family. Finally, Sncg t-types were divided into two classes according to Vip expression, with Sncg-Vip and Sncg-Pdzrn3 identified as small and large Cck cells, respectively^113,114^.

### Data analysis

#### Modulation Index

When comparing activity in two conditions (e.g. visual stimulus vs. blank; large vs. small grating; running vs. Stationary synchronized), we used a modulation index computed as

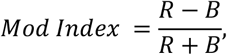

where R the mean activity during the response time window and B the mean activity during the baseline time window:

#### Cell depth comparison to Patch-Seq study

For the analysis validating coppaFISH t-type calling using cell depth (Fig. 1k), we used cells of all layers, not just the *in vivo* imaged cells of L1-3. We used 14 sections for which gene expression was obtained from layer 1 to layer 6 (all taken from the same animal). DAPI segmentation was manually curated (see above) in all layers, and cell calling was performed on these sections using the standard method. This provided the cortical depth for about 47000 cells among which 2130 were assigned to a GABAergic t-type. We normalized the measured cortical depth by the maximum cortical depth in these sections (750 microns) and computed the median cortical depth for each t-type with at least 4 cells (46 such t-types were found). We then did the same thing for the Patch-seq data of Gouwens et al.^8^, which gave 42 t-types with more than 4 cells. We then compared the cortical depth of the t-types with at least 4 cells in both datasets (33 t-types in total; Fig. 1k).

#### Determining behavioural states

To distinguish the 3 main behavioural states during spontaneous behaviour, we used the running speed of the animal as well as the strength of cortical oscillations. Running speed was measured by optical sensors facing the air-suspended ball^115^, and was smoothed with a 2 s moving average filter. We considered the mouse stationary if this smoothed speed was less than 0.3 cm/s, and running otherwise. To distinguish between the synchronized and desynchronized stationary states, we first computed the first principal component of excitatory cells’ activity using PCA, which revealed cells more active in passive or alert states, as previously described^49^. The activity of the 10% of cells with highest weight on this PC was averaged, which provided a clear summary of the oscillation that appeared in some stationary periods (Fig. 2a). Periods of synchronized activity were segmented manually based on the periods where this average was clearly oscillating. To measure the oscillatory coupling of each inhibitory neuron, we then computed the correlation between each cell’s z-scored activity and the average of this excitatory subpopulation during the synchronized periods.

#### Comparison to transgenic mouse line data

To validate our cell type assignment, we compared results obtained with post-hoc transcriptomic with recordings performed using transgenic mouse lines (Extended Data Fig. 5). We analysed recordings from 18 transgenic mice (5 for Pvalb, 8 for Sst, and 5 for Vip; 14 mice were reanalysed from Ref. ^19^ and 4 new mice were added) and 23 sessions (6 for Pvalb, 9 for Sst, and 8 for Vip) for a total of 2,589 identified cells (1023 Pvalb, 572 Sst and 994 Vip cells).

For this analysis (Extended Data Fig. 5), we first deconvolved the calcium traces to inferred firing rates (*t*) for each neuron *i* at time *t*^84^. We considered two measures of neural activity for each cell *i* and trial *n:* the average neural activity *r_i_*(*n*) = 〈*f_i_*(*t*)〉_*tϵ*_[*t_n_,t_n_*+Δ*T*] during stimulus presentation from the trial onset time *t_n_* to time *t_n_* + Δ*T*, and the average neural response *d_i_* (*n*) = *r_i_*(*n*) – *b_i_*(*n*), obtained after subtracting the pre-stimulus baseline activity *b_i_*(*n*) = 〈*f_i_*(*t*)〉_*tϵ*_[*t_n_*–Δ*T,t_n_*]. The time window parameter Δ*T* took the value 1 s for the data from Ref.^19^ and 0.5 s for the new transgenic data and the post-hoc transcriptomic data, corresponding to the whole duration of the stimulus. We then computed the average activity and response for a given stimulus s and locomotion condition 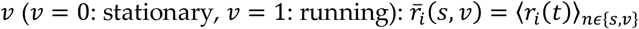 and 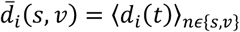.. We estimated the responsiveness of each neuron *i* to visual stimuli by computing the p-value *p*, of a paired t-test comparing η (*n*) with *b_i_*(*n*) for all trials *n* (pooling all different stimulus types to obtain one p-value per cell). For all subsequent analysis we selected only cells with p-values < 0.05. We plotted the average modulation of visual responses by running (computed as in Ref.^19^) vs. the Pearson correlation coefficient of spontaneous activity and running speed *ρ_i_* (Extended Fig. 5a). Prior to computing the Pearson correlation coefficient, we smoothed the activity *f_i_*(*t*) and running speed *v*(*t*) with a time average of 5 s. For this analysis, we selected only cells whose cortical depth was > −300 *μm*.

For estimating size tuning curves (Extended Data Fig. 5b), we z-scored the activity of each neuron as follows 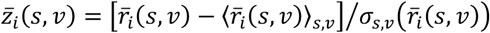 prior to averaging over cells of a given type.

To evaluate consistency between the physiological features identified with transgenic and transcriptomic cell type identification, we trained a classifier to predict cell type from physiological features of each cell in the transgenic lines, and asked if it generalized to the transcriptomic data (Extended Data Fig. 5c). We trained the classifier using 1,230 training cells (410 examples per cell type for the three cell types). The prediction was based on 14 features, which included normalized values of neural activity during different stimulus size and running condition 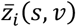 (features #1-8); skewness of the calcium trace computed across the whole recording session (feature #9); the correlation of spontaneous activity with running speed *ρ_i_* (feature #10); the ROI diameter (feature #11); the cortical depth (feature #12); two different measures of the difference in modulation by running 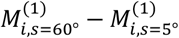 between large and small stimuli, where 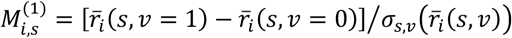 and 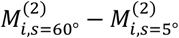, where 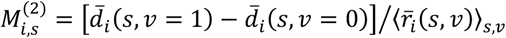 (features #13 and #14). We normalized features #9-14 by z-scoring them using the mean and standard deviation for each neuron of the transgenic mice, while features #1-8 were already normalized as in Extended Data Fig. 5b. We used cell types: *y =* {F% *SOM, VIP]* as training labels. Using the 10 different randomized splits of training and test transgenic data, we applied three different linear classifiers: Linear Discriminant Analysis, Logistic Regression (regularization parameter C = 10) and Linear Support Vector Classification (regularization parameters C = 0.1). The regularization parameters were chosen after a 4-fold cross-validation over the different randomized training sets scanning over *C* = {10^-3^≡,10^-2^, …, 10^2^}. Applying the classifier to transcriptomic data gave equivalent performance to test-set transgenic data, indicating that the two methods are consistent.

#### Response to drifting gratings

Responsive cells (either Activated or Suppressed) were defined using a repeated measures ANOVA model (fitrm in Matlab) with the stimulus direction (12 levels) and size (3 levels) as between subjects factors, and the presence of stimulus as a within subject factor. A cell was defined as responsive if there was a significant effect of stimulus presence after performing a repeated measures analysis of variance (ranova in Matlab). Significant cells were classified as Activated if mean activity in the response window was above baseline, or Suppressed otherwise.

Orientation selectivity Index (OSI) was computed using a cross-validation method. Each cell’s preferred orientation was computed from even trials, selectivity was computed as:

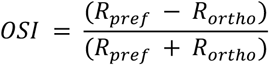

Where R_pref_ is the mean response on the odd trials to the preferred orientation and R_ortho_ is the mean response on the odd trials to the orthogonal orientation (R_pref_ + 90°). This cross-validation was used because non-cross-validated selectivity indices can show large values for sparse neural activity, even if the cells are untuned. The cross-validated measure can take negative values, which indicate inconsistent responses, and will have an expected value of 0 for untuned cells.

Direction selectivity Index (DSI) was obtained similarly. Each cell’s preferred direction was computed from even trials, selectivity was computed as:

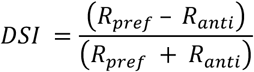

Where R_pref_ is the mean response on the odd trials to the preferred direction and R_anti_ is the mean response on the odd trials to the direction opposite to the preferred (R_pref_ + 180°).

Size tuning curves and their state modulation (Fig. 3e) were computed using the methods of Ref.^19^. Analysis was restricted to cells whose receptive field locations were close to the centre of the grating stimuli (<20°). Size tuning curves were obtained for running and stationary states by averaging the z-scored activity of all centred cells of that class (z-scoring was computed relative to the entire recording session). Baseline activity (shown as response to size 0 stimuli) was estimated as the average of the z-scored activity during the interstimulus intervals. For both the stimulus response and the baseline, we determined if the mouse was running or stationary by taking the average running speed during the stimulus presentation. If this speed exceeded 1cm/s we considered the mouse as running, and stationary otherwise.

Cross-validated direction tuning curves (Fig. 3b) were computed for all cells using the average across all sizes. A cell’s preferred direction was estimated as the direction providing the largest response on even trials. Direction tuning curves were computed by averaging the z-scored activity of each cell on odd trials, for each direction relative to this preferred direction. The curve was normalized by dividing by the mean response to the preferred direction (on the even trials). These normalized curves were then averaged over all cells in a t-type (Fig. 3b).

#### Pairwise correlations between Classes

To compute spontaneous correlations between the mean activity of Class (Fig. 4d), we first normalized each cell’s deconvolved activity by dividing it by its maximum. For each experiment, we then averaged the normalized activity of each cell within a Class during blank-screen periods, smoothed with a 1s boxcar window, and decimated the sampling rate to 1 Hz. We computed the Pearson correlation between each Class’s mean activity and averaged over experiments. For the intra-Class correlations, we randomly split the cells of each Class in two halves and applied the same method, to avoid trivially obtaining a correlation of 1. When the number of cells in a Class was less than 4, the correlation was not computed for that experiment.

#### Response to Natural Images

We summarized a cell’s response to natural image stimuli with two numbers (Fig. 3d). Responsiveness was defined as a modulation index between activity during the stimulus presentation period and the activity just before stimulus onset. Signal correlation was defined by correlating the responses to the first repeat of the 1050 images with the responses to the second repeat of these same images. This metric characterizes a cell’s selectivity to these image stimuli^116^.

#### Genetic PCA

To compute the first genetic principal component, we averaged the *in situ* gene expression of the 72 genes for each of the 35 t-types. We then performed PCA on this 72 by 35 matrix, and took the score of the first component to get gPC1 for each t-type. To obtain gPC1 values for cells in Patch-seq (Extended Data Fig. 9), the same weight vector was used and read counts were transformed by log(1+x).

#### UMAP on Tasic et al scRNA-seq data

We performed a UMAP analysis on the Tasic et al scRNA-seq dataset^4^, separately for CGE (Vip, Sncg and Lamp5) and MGE (Pvalb and Sst) derived inhibitory Families from V1 only (Extended Data Fig. 3).

To do so, we employed methods previously described for CA1^13^. First, a set of 150 genes was found using the ProMMT clustering algorithm. 150-dimensional expression vectors were made for each cell, applying a log(2+x) transform to the scRNA-seq expression levels of these genes. UMAP analysis was performed using Meehan et al’s Matlab toolbox^117^, initialized by placing the classes around a unit circle in order of similarity.

The genes automatically selected to perform the UMAP analysis were: Vip, Tac2, Sst, Pdyn, Lamp5, Tac1, Crh, Calb1, Penk, Calb2, Th, Cxcl14, Ndnf, Spp1, Htr3a, Cplx3, Pvalb, Crhbp, Npy, Npy2r, Chodl, Crispld2, Prss23, Nov, Cbln2, Cartpt, Akr1c18, Atp6ap1l, Cadps2, Ppapdc1a, Sncg, Tnfaip8l3, Unc13c, Pdlim3, Scgn, Pcp4, Tcap, Lgals1, Serpine2, Moxd1, Pthlh, Cd34, Cck, Sostdc1, Spon1, LOC105243425, Mia, Slc5a7, Pde1a, Adarb2, Mybpc1, Car4, Cbln4, Gabrg1, Fmo1, Slc18a3, Grpr, Lypd6, Pde11a, Rxfp1, Tnnt1, Nxph2, Lpl, Cryab, Cp, Npy1r, Id3, Myl1, Id2, Kit, Serpinf1, Bcar3, Aqp5, Scrg1, Gpd1, Rxfp3, Prox1, Col25a1, Chat, Vwc2l, Amigo2, Myh8, Synpr, Grm8, Igfbp5, Gpx3, Rgs12, Lypd1, Cd24a, Reln, Hapln1, Sln, Chrm2, Ostn, Igfbp7, LOC102632463, Atf3, Lect1, Gpc3, Ptprk, Teddm3, Il1rapl2, Col6a1, Nek7, Crispld1, Wif1, Wnt5a, Bmp3, Thrsp, Syt2, Pcdh20, Sfrp2, Myh13, Efemp1, Rprm, Cacna2d1, Lypd6b, Meis2, Lhx6, Angpt1, Rspo1, Sema3c, Itih5, Nfix, Sema3a, Stk32a, Ecel1, Jam2, Igfbp6, Sox6, Nfib, Sall1, Sema5b, Shisa8, Tacr3, Chst7, Frmd7, Gm31465, Rspo4, Chrna2, Lmo1, C1qtnf7, Ndst4, Ccdc109b, Npas1, Egfr, S100a10, Gpr6, Slit2, Lsp1.

#### Correlation with electrophysiological and morphological properties

We examined electrophysiological and morphological correlates of our results by relating them to a previously published Patch-seq dataset^8^, which provided electrophysiological, morphological, and gene expression data from a set of V1 inhibitory cells analysed *in vitro.* These cells had been genetically assigned to the same transcriptomic clusters we used^4^, which allowed us to correlate electrophysiological and morphological properties to the state modulation measured in our own dataset. Valid electrophysiological recordings were available for 4391 cells and included long and short pulses of current injection as well as current ramps. We used the electrophysiological parameters calculated by the original authors using the ipfx software, renaming “up/down ratio” (the absolute ratio of the slopes of the upward and downward components of the action potential) as “spike shape index”. Adaptation index was the rate at which spiking changed during a long depolarizing square stimulus. During a hyperpolarizing square current, the membrane time constant tau is the rate of approach of steady state, and sag is the downward deflection before steady state is reached. Capacitance was calculated as the ratio between measured tau and resistance.

We quantified the ratio of axon in each layer using morphological reconstructions obtained following Patch-seq. To enable comparison to our 2-photon data, we only examined reconstructed cells with somas in layers 1-3 that belonged to one of the 35 t-types we recorded from, for a total of 163 cells. Morphology was represented as an acyclic undirected graph with a position and radius associated with each node. A pair of adjacent nodes (a segment) fell within a layer if both nodes had cortical depths within the layer boundary. Segments which fell on a layer boundary (less than 4% of segments for each cell) were not classified into a layer, and segments entering the white matter or pia were excluded. The surface area of all within-layer segments was computed using the distance between nodes and their radii. The within-layer surface area ratio is the sum of the surface area of segments within a layer divided by the total surface area of all segments.

gPC1 was computed for each Patch-seq cell using the same 72 genes and weightings found from our coppaFISH data, with gene expression transformed as log(1+x).

#### Processing of eye video (pupil detection)

Eye videos were processed using facemap (https://github.com/MouseLand/facemap). An ROI was drawn manually around the pupil of the animal. The pupil area was defined as the area of a Gaussian fit on thresholded pupil frames, where pixels outside the pupil were set to zero.

## Statistical analyses

Statistical analysis of differences between cell types faces two potential confounds. First, different experiments will by chance record different proportions of each cell type, and may also by chance show other experiment-to-experiment differences such as overall alertness levels. Second, the large number of t-types presents a potential multiple comparisons problem.

To solve these problems, we used a hierarchical permutation test. First, an Omnibus test asks whether Family, Class, and t-type have a significant main effect on our quantity of interest *y*; there is no multiple-comparisons problem for this Omnibus test, and all shuffling occurs within an experiment to avoid conflating experiment-to-experiment variability with differences between cell types. The Omnibus test is conducted at each of the 3 levels in a nested manner: the first asks if there is a main effect of Family; the second if there is a main effect of Class beyond that predicted by Family; and the third if there is a main effect t-type beyond that predicted by Class. Following the Omnibus test, post-hoc tests are used to ask if significant differences between Classes exist within each individual Family, and if significant differences between t-types exist within each individual Class. Additional post-hoc tests are used to ask whether the quantity is significantly different to zero for each Class and t-type. All post-hoc tests are corrected for multiple comparisons using the Benjamini-Hochberg procedure.

To test for a main effect of Family on a quantity *y*, the Omnibus test computes its mean value of 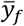 for each family *f*, and uses as test statistic the variance of 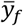 across families. To obtain a p-value, this test statistic is compared to a null ensemble obtained after 10,000 random shufflings of the Family label of each cell, separately within each experiment. To test for a main effect of Class, we compute the mean 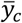 of *y* for each Class *c*, and use as test static the variance of this mean across Classes. A null distribution is obtained by 10,000 shufflings of Class labels separately within each experiment and Family. To test for a main effect of t-type, we use as test statistic the variance of 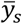 over t-types *s*. A null distribution is obtained by recomputing this statistic after shuffling t-type labels 10,000 times, separately within each Class and experiment.

To perform the post-hoc test for significant differences between the Classes within a specific Family (indicated by p values on the far right of Fig. 2b and similar), or for significant differences between t-types within a specific Class (indicated by stars second to right in Fig. 2b), we performed the same shuffle test inside individual Families and Classes. For example, to obtain the p-value for significant differences of t-types within the Pvalb-Tac1 Class, we used as test statistic the variance of 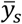 across the 5 t-types inside this Class, and compared it to 10,000 shufflings of the t-type labels inside this same Class. These post-hoc p-values were then corrected using the Benjamini-Hochberg procedure. For post-hoc tests of whether a Class or t-type is significantly different to zero, we used Benjamini-Hochberg corrected t-tests.

For linear correlations (Fig. 1k, Fig. 2e, Fig. 4e-f Extended Data Fig. 6c, Extended Data Fig. 9a-b), we show the p-value for the Pearson correlation coefficient. To exclude the possibility of conflating experiment-to-experiment variability with differences between cell types, we used ANCOVA controlling for a discrete effect of recording session; (Fig. 2c-e, Fig. 4b-c, Extended Data Fig. 6c,) quoting the significance of a main effect of the continuous variable. For Fig. 2c-d and Fig. 4b-c, we performed the ANCOVA after averaging the relevant values of all cells for a given recording session and t-type. ANCOVA was also used to test whether a continuous genetic variable assigned to each cell correlated significantly with state modulation even after controlling for t-type and recording session (Fig. 2e, Extended Data Fig. 6f), and if cortical depths of each t-type measured by coppaFISH and Patch-seq were correlated even within a Family or Class (Fig. 1k).

To test for the effect of gPC1 on pairwise correlations (Fig. 4d), we sorted Classes by gPC1 and computed their pairwise correlation matrix as described above. We used a permutation test to ask if values close to the diagonal were larger than values far from the diagonal. As test statistic we used the difference between the mean correlation values one or two steps away from the diagonal, and the mean of all other class pairs (Extended Data Fig. 9d). We constructed a null distribution by recomputing this statistic after permuting the order of the Classes 10,000 times.

## Acknowledgements

We thank Thomas Hauling for work developing the coppaFISH method; Mats Nilsson and Xiaoyan Qian for advice with *in situ* transcriptomics; Nathan Gouwens for help with Patch-seq data; Federico Rossi for discussion; Michael Krumin for support with microscopy; Laura Funnell for help with histology. This work was supported by the Wellcome Trust (108726, 205093, 090843), the European Union’s Marie Skłodowska-Curie program (835489), the Royal Society (NIF\R1\ 180184) the Chan-Zuckerberg initiative (2018-182811), and the Gatsby Charitable Foundation (GAT3361). MC holds the GlaxoSmithKline / Fight for Sight Chair in Visual Neuroscience.

## Author contributions

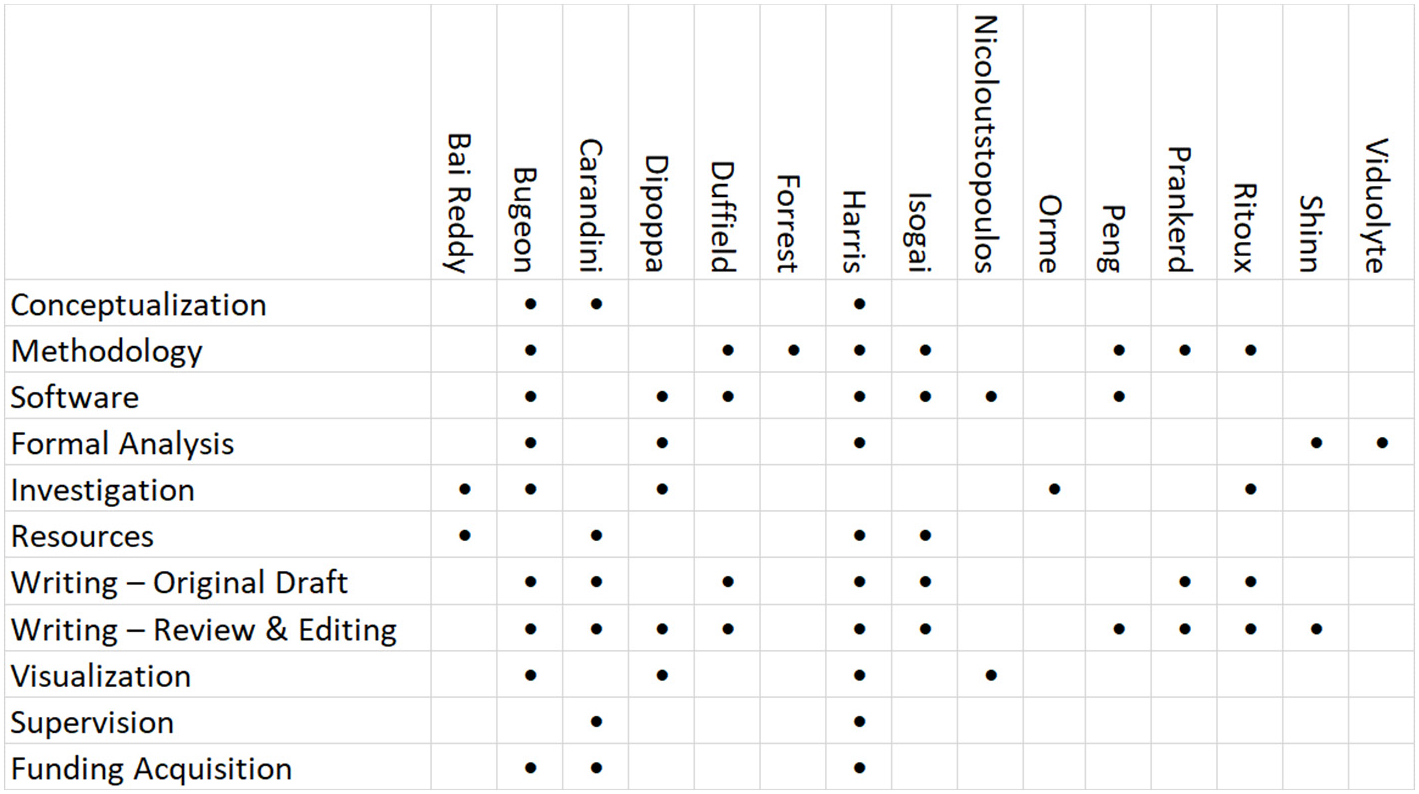

## Extended Data Figures

**Extended Data Figure 1.**
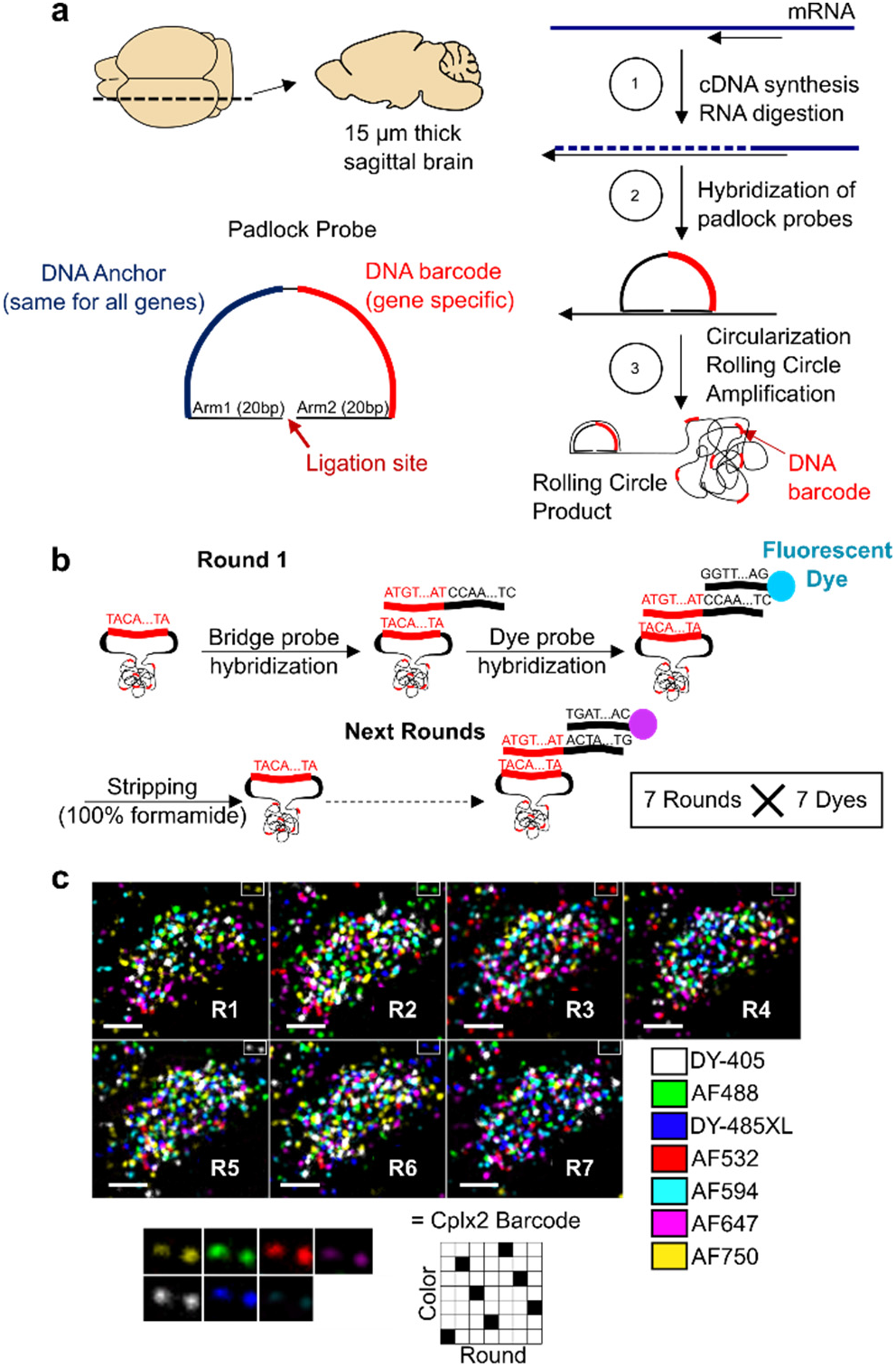
Detection of 72 genes using coppaFISH. **a**, Sagittal 15 μm brain sections are cut using a cryostat. Local mRNAs are retro-transcribed to cDNA, and the mRNAs digested to free the cDNAs for hybridization with padlock probes. Padlock probes have two 15-20 nucleotide (nt) arms complementary to the target site, a 20nt anchor sequence (identical for all probes) and a 20nt barcode sequence (unique for each gene). After hybridization to the target site, a DNA ligase enzyme circularizes the padlock probe, but only when it matches the target perfectly. Next, a DNA polymerase enzyme amplifies the circularized padlock probes, producing rolling circle products (RCPs), which contain many repeats of the padlock sequence including the barcode. **b**, The genes are detected by 7 rounds of 7-colour fluorescence imaging. On each round, RCPs are hybridized with custom designed bridge probes, which in turn hybridize to specific dye probes (conjugated to one of 7 fluorophores). The sections are then imaged in 7 colour channels, then all DNA is removed with formamide treatment, and the next round begins. Different sets of bridge probes on each round result in each barcode showing up in a different colour channel using a Reed-Solomon code for minimum overlap. After the 7 combinatorial rounds, a final round images the anchor probe (used for image alignment) and DAPI to visualize cell nuclei. **c**, Example raw data for one cell imaged with the 7 fluorophores and 7 rounds. Each fluorescent spot is an RCP, and the sequence of colours across 7 rounds allows gene identity to be determined. Bottom: magnification of 2 RCPs (top right corner of main images) which corresponded to Cplx2 barcode (6135024). Scale bars: 5 μm.

**Extended Data Figure 2.**
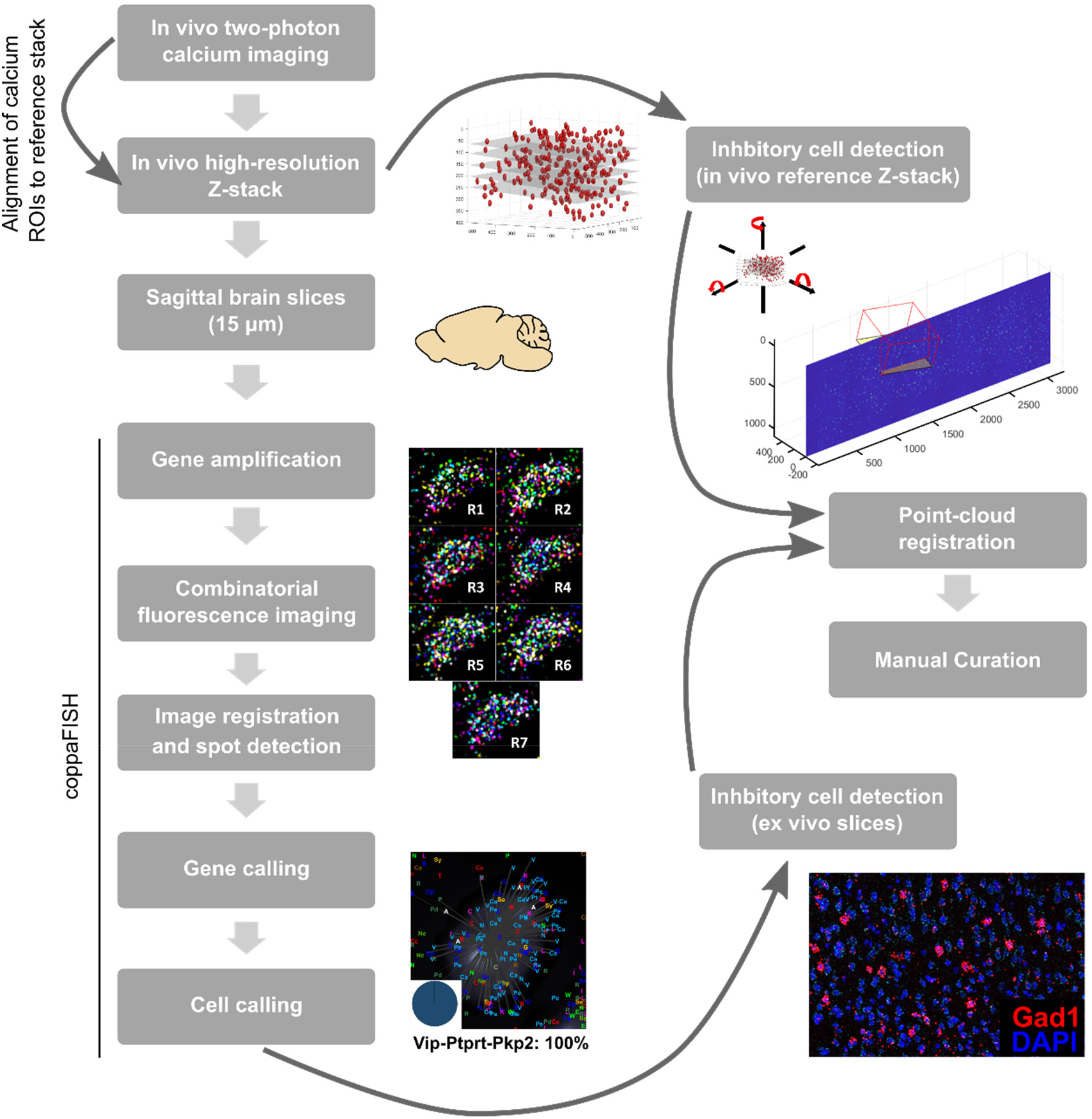
Experimental pipeline. Neural activity was recorded *in vivo* over multiple sessions from each subject (Gad2-mCherry mice with viral GCaMP expression in all neurons). At the end of each session, a high-resolution reference Z-Stack was acquired and used to detect interneurons in the Z-stack volume using mCherry fluorescence, and cells recorded during calcium imaging were registered to this Z-Stack. After all imaging sessions, the brain was extracted from the skull without fixation and frozen in OCT. A block from under the imaging window was sliced into 15 μm sagittal sections, which were thaw-mounted on gelatine-coated coverslips. Each section was then processed using coppaFISH: RCPs were produced *in situ* for the selected genes, and their barcodes were read using 7 rounds of imaging (+ 1 anchor round). The resulting images were then registered across rounds, colour channels, and image tiles and individual spots detected. Gene identity for each RCP was decoded from the 49-dimensional images, and pciSeq^41^ was used to determine the t-type identity for each cell. To align the images, interneurons detected *in vivo* and *ex vivo* were used as fiducial markers for point cloud registration, which finds the best alignment of the 2D *ex vivo* slice in the 3D volume. Finally, individual cell matches were manually curated, and a t-type assigned to the recorded cells.

**Extended Data Figure 3.**
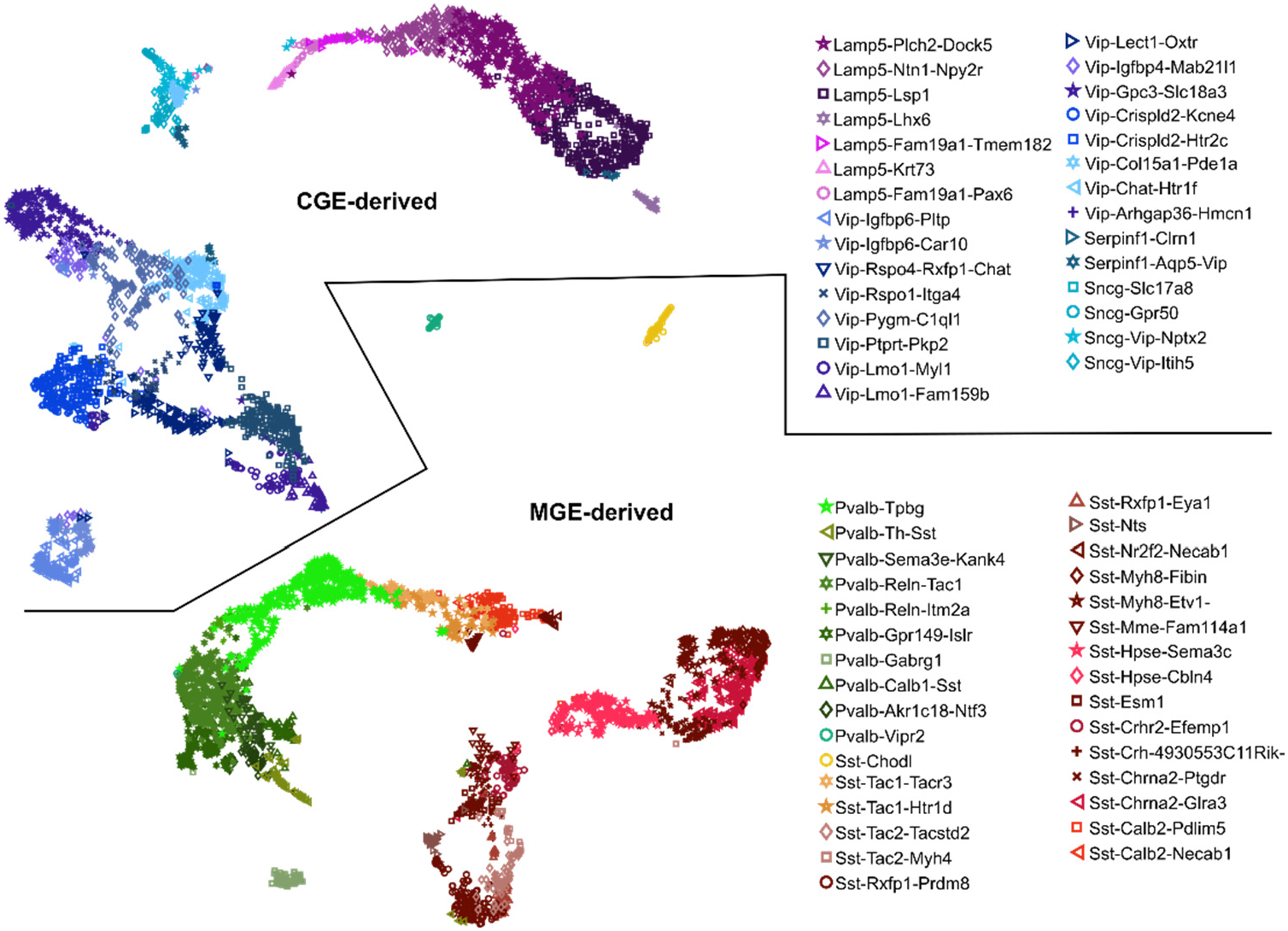
UMAP analysis of scRNA-seq data. Each dot represents a V1 inhibitory cell, from the Tasic et al.^4^ data, with glyph representing its assigned t-type. UMAP analysis was performed separately for MGE and CGE derived interneuron t-types, using 150 log-transformed genes selected by the ProMMT algorithm^13^. This analysis reveals both highly discrete t-types such as Pvalb-Vipr2 (putative chandelier cells) and smoothly varying continua where boundaries between t-types appear arbitrary, such as Lamp5-Ntn-Npy2r, Lamp5-Plch2-Dock5, and Lamp5-Lsp1 (putative neurogliaform t-types). Also note the smooth transition between Sst-Calb2 (a putative Martinotti subtype), Sst-Tac1 (putative Sst non-Martinotti), and Pvalb-Tpbg (putative superficial basket) cell t-types.

**Extended Data Figure 4.**
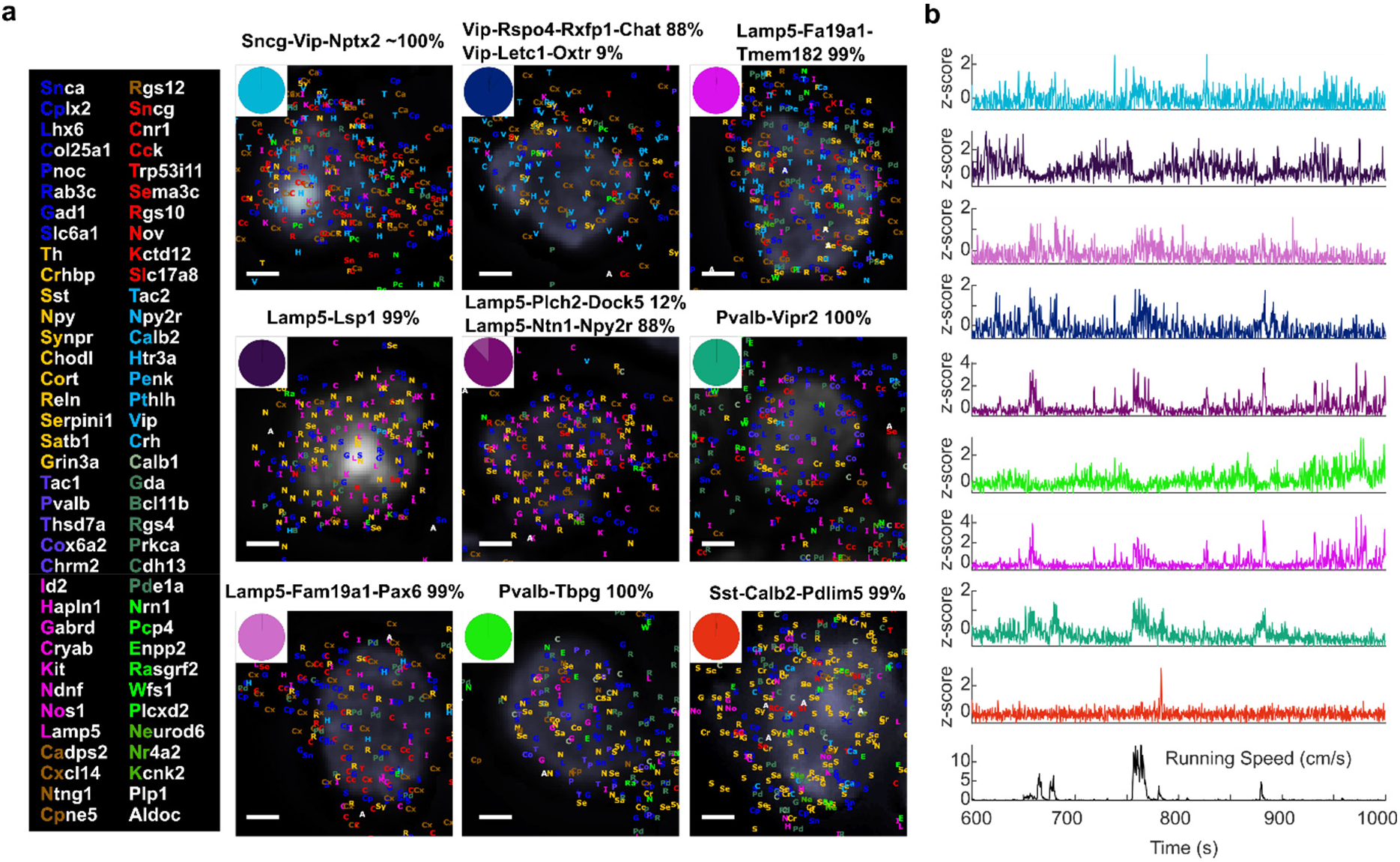
Example Cells. **a**, Nine example cells which were recorded during the same session. Pie plots indicate the posterior probability of each cell’s t-type assignment. Grey background images show DAPI-stained nuclei. Each gene detection is represented by coloured letters (key to the left). Scale bars: 2 μm. **b**, Activity of these 9 cells during spontaneous behaviour, together with the running speed of the animal. The traces are colour coded according to the assigned t-type for each cell (pie plots in **a**).

**Extended Data Figure 5.**
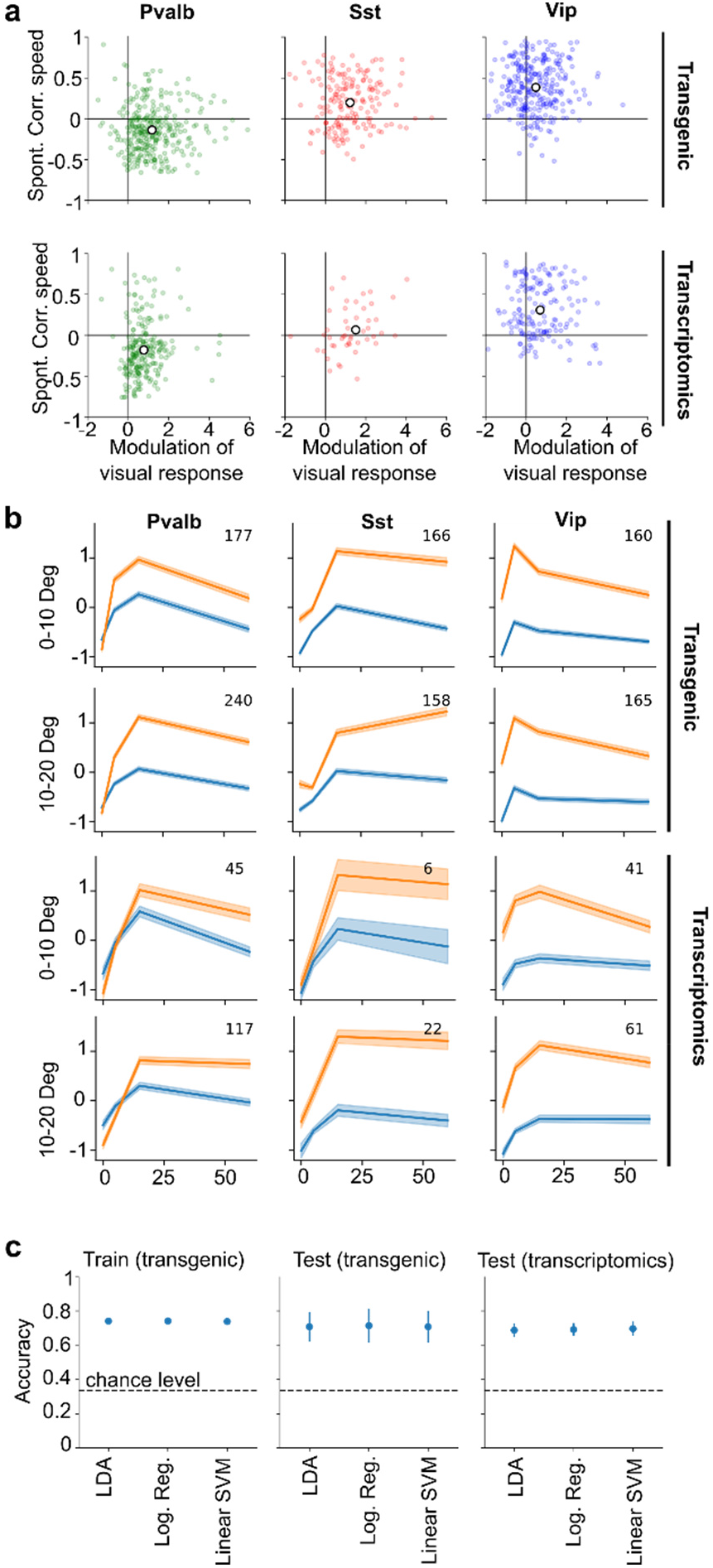
Comparison to results in transgenic mice. **a**, Top row: modulation of visual responses by running vs. correlation to running speed during spontaneous behaviour, for Pvalb, Sst, and Vip interneurons identified in transgenic mouse lines. Data re-analysed from Ref.^19^ and including 4 new animals. Bottom row: same analysis using interneurons identified by post-hoc transcriptomic analysis (data from this study; the Vip group included Vip-positive Sncg cells which are likely to be labelled in the Vip-Cre transgenic line). In both datasets, running suppressed the spontaneous activity of Pvalb cells, but enhanced their visual response. In both datasets, Sst cells showed weakly positive spontaneous correlation to running and stronger positive modulation of visual responses. In both datasets, Vip cells showed stronger modulation by running during spontaneous behaviour than during visual stimulation. **b**, Size tuning curves of Vip, Pvalb and Sst cells for both datasets. Top row: responses measured in transgenic mice for centred stimuli (0-10° offset from receptive field centre); second row: response to off-centre stimuli (10-20° offset from receptive field) in transgenic mice; bottom two rows, same from post-hoc transcriptomics. Orange curves: responses during running; blue curves, responses during stationary epochs. Note that in both cases, Vip cells responded more to small than large stimuli; Sst cells showed little surround suppression by large stimuli and responded weakly to small stimuli; and Pvalb cells showed an interaction of stimulus size and behaviour, with larger running modulation for larger stimuli. **c**, Classification of cell type from physiological features was identical for the two cell typing methods. Each cell was assigned to either Sst, Pvalb or Vip based on 14 physiological features (such as correlation to running speed, size tuning curves, skewness), using one of 3 different linear classifiers trained on a training set randomly selected from the transgenic recording sessions. Left: training-set classifier accuracy averaged over multiple random selections of the training set. Centre: accuracy of the classifiers on the held-out transgenic sessions, averaged over randomized splits into training and test sessions. Right: out-of-sample accuracy of the linear models on data with interneurons identified by post-hoc transcriptomics. Note the similar performance on transgenic and transcriptomic test sets. Error bars: s.d. over divisions into training and test set.

**Extended Data Figure 6.**
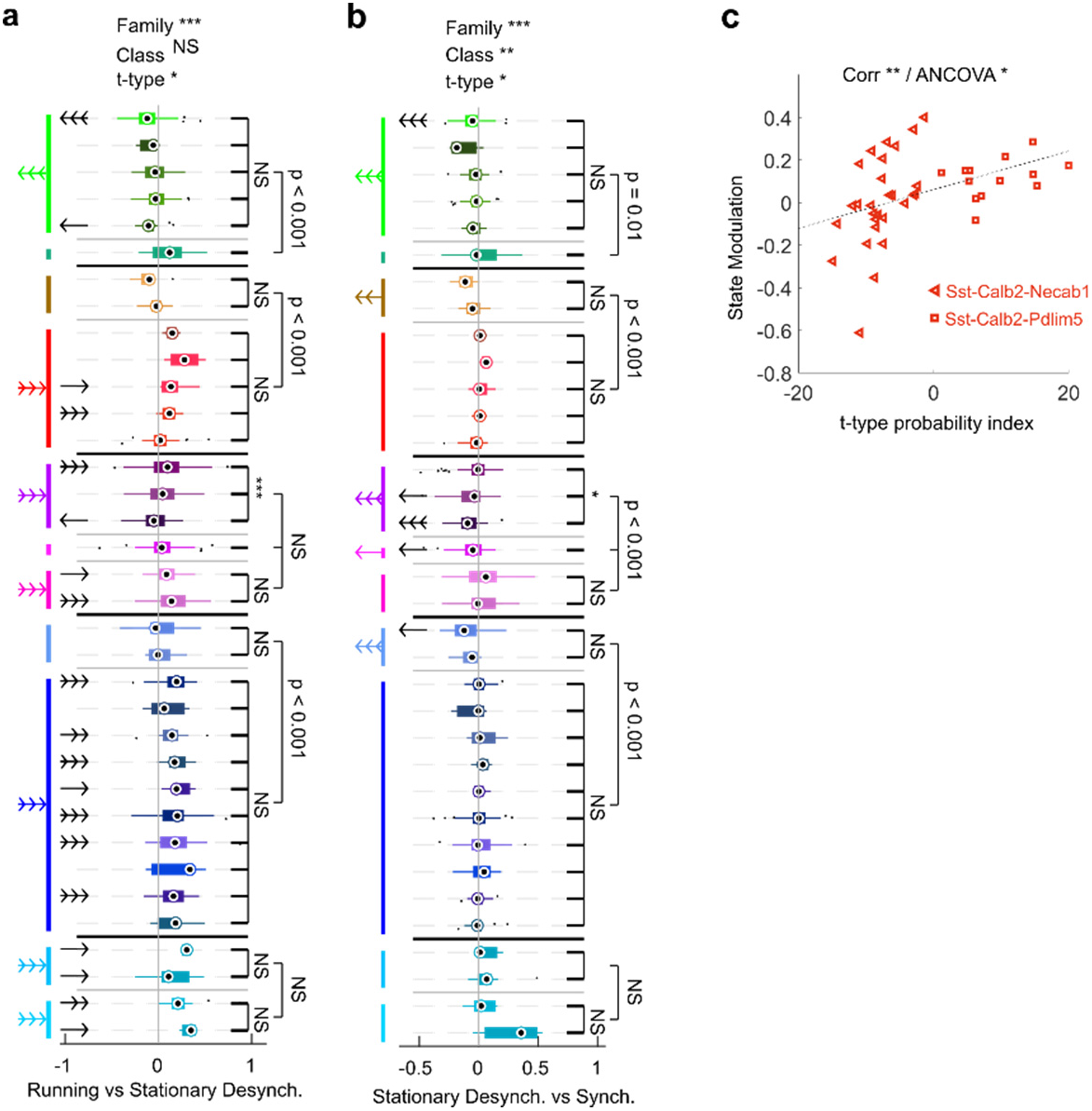
Further analyses of state modulation in spontaneous behaviour. **a**, Hierarchical analysis of modulation between Running state and Stationary Desynchronized state, plotted as in Fig. 2b. **b**, Hierarchical analysis of modulation between Stationary Desynchronized and Stationary Synchronized states, plotted as in Fig. 2b. **c**, State modulation vs. t-type probability index for Sst-Calb2-Necab1 and Sst-Calb2-Pdlim5 cells (p<0.01, Pearson correlation; p=0.013, ANCOVA accounting for effects of session and t-type).

**Extended Data Figure 7.**
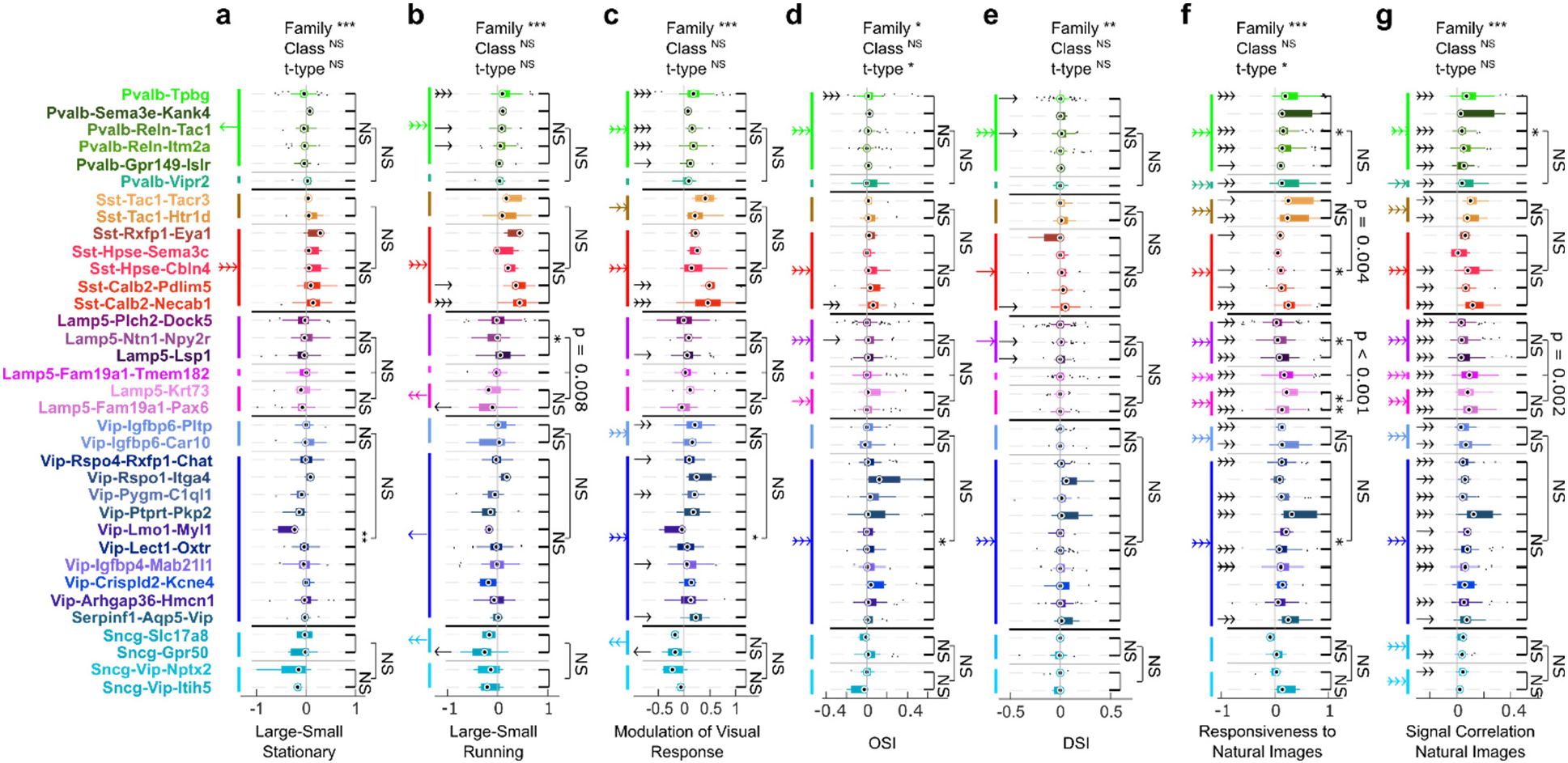
Further analyses of visual responses. Each panel shows a hierarchical analysis for the visual variables analysed in Fig. 3d, but showing all t-types. All panels plotted as in Fig. 2b.

**Extended Data Figure 8.**
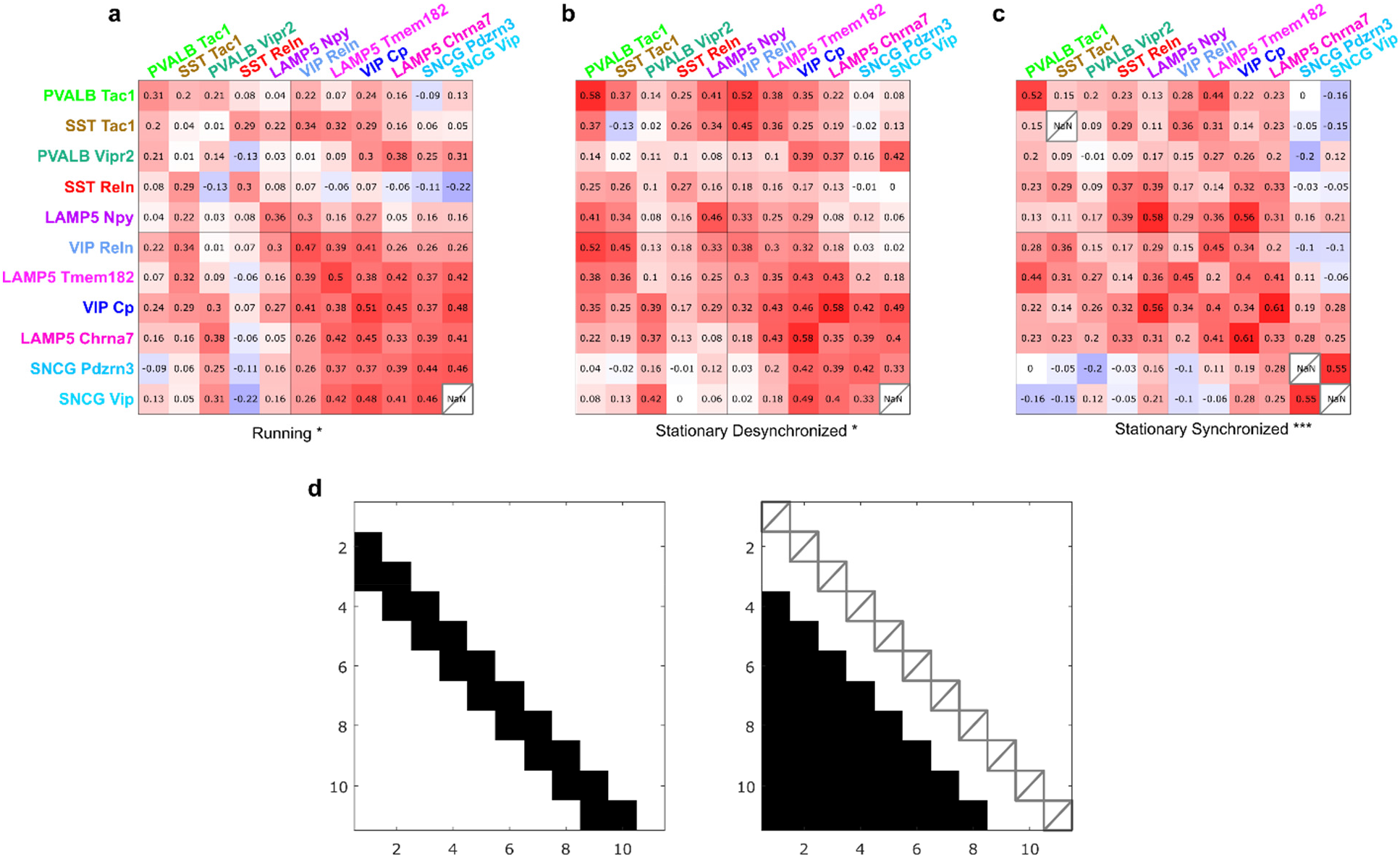
Analysis of pairwise correlations within states. **a**, **b**, **c**, Pairwise correlations between simultaneously recorded Classes, plotted as in Fig. 4d, but separately for periods within each of the three states (Running, Stationary Desynchronized, and Stationary Synchronized). The Classes are sorted by gPC1; Classes with similar gPC1 values have significantly higher correlations (permutation test, p=0.018, p=0.037, p=0.0008 respectively). **d**, The test statistic for the permutation test was the difference between the average of correlation coefficients close to the diagonal (left), and the average of all other off-diagonal coefficients; intra-class correlations were not used. This test statistic was compared to a null ensemble obtained after shuffling gPC1 values 10,000 times.

**Extended Data Figure 9.**
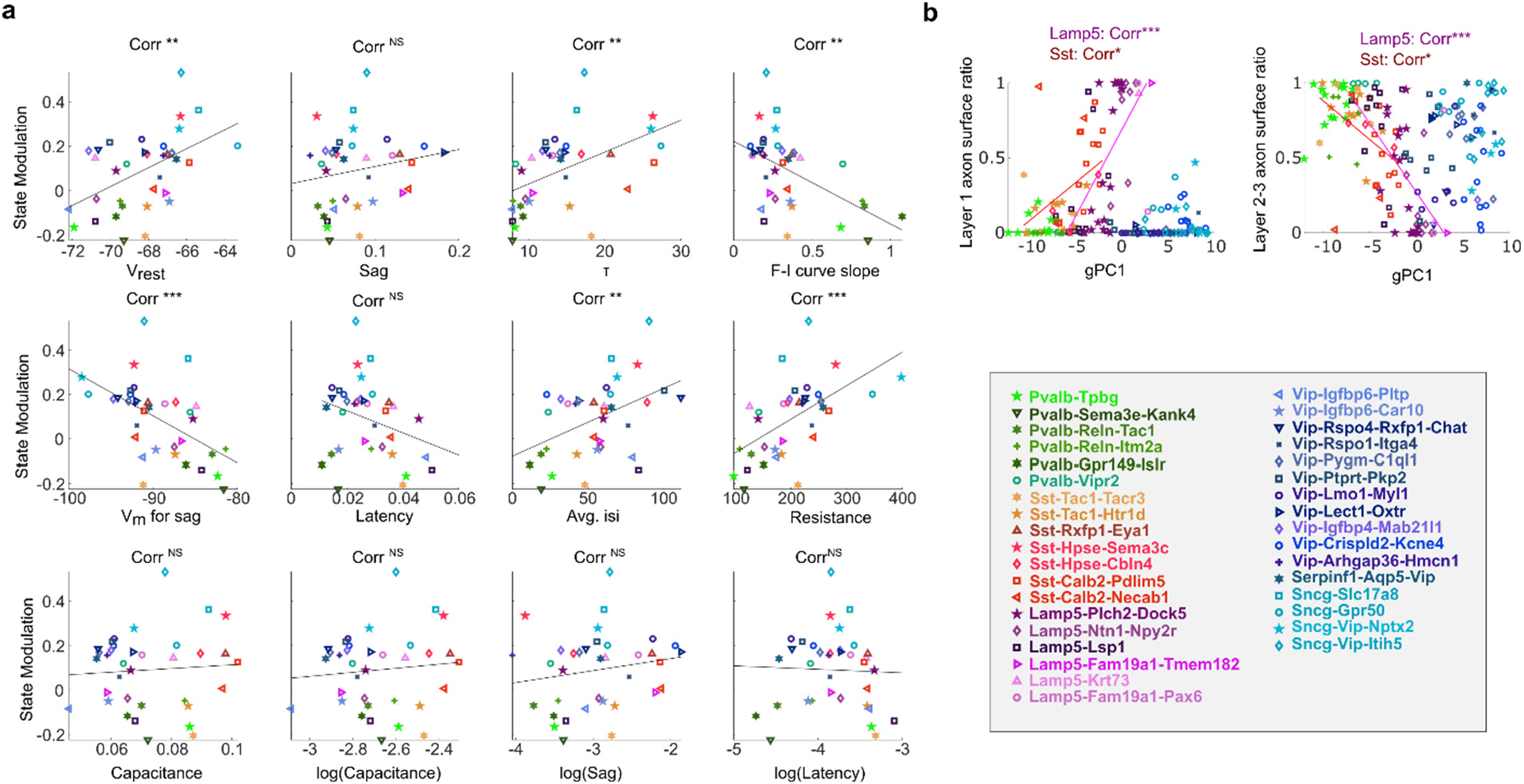
Additional analyses of Patch-seq data. **a**, Additional electrophysiological properties vs. State modulation plotted as in Fig. 4e. V_rest_: r=0.25, Sag: r=0.03, τ: r=0.24, F-I curve slope: r=0.28, V_m_ for Sag: r=0.29, Latency: r=0.09, Avg. isi (inter-spike interval): r=0.20, Resistance: r=0.34, Capacitance: r=0.01, log(Capacitance) : r=0.01, log(Sag) : r=0.03 and log(Latency): r=0. Stars show significance assessed by Pearson correlation. Dashed lines are linear fits. **b**, Fraction of axonal arborization (measured by surface area) in layer 1 (left) and layer 2-3 (right) vs. gPC1 computed for each Patch-seq neuron. Each symbol represents a cell. Pearson correlation was computed individually within each Family, and p-values were adjusted with Benjamini-Hochberg correction (Layer 1 Lamp5: r=0.40; Layer 1 Sst: r=0.17; Layer 2-3 Lamp5: r=0.34; Layer 2-3 Sst: r=0.19). Coloured lines show linear fit for each Family with significant Pearson correlation. *, p<0.05, **, p<0.01, ***, p<0.001.

**Supplementary Data File 1. Percentage of interneurons assigned to a t-type.** Number of interneurons recorded per session and per animal, and percentage of interneurons that were assigned to a t-type at the end of the experimental pipeline. In total, about 44% of recorded interneurons were characterized transcriptomically

**Supplementary Data File 2. Padlock probe sequences.** Name and sequence of the 556 padlock probes (73 to 80nt) targeting the cDNA sequences produced by reverse transcription. Each probe contains the same 20nt anchor sequence, a 20nt gene specific DNA barcode, and two arms complementary to the cDNA sequence.

**Supplementary Data File 3. Primer sequences**. Name and sequence of the 556 primers used for reverse transcription of the mRNAs.

**Supplementary Data File 4. Dye probe sequences.** Name and sequence of the 7 dye probes used for combinatorial imaging. Each 20nt DNA oligo was conjugated to a given dye. All dyes were conjugated at the 5’ end only, except from dp0 and dp6 which were conjugated at both ends.

**Supplementary Data File 5. Bridge probe sequences.** Name and sequence of the 511 bridge probes used for combinatorial imaging, 1 for each gene and imaging round.

**Supplementary Data File 6. Reed-Solomon codes.** Dye code for each gene, consisting of 7 numbers between 0 and 6, listing the dye probe that gene is assigned on each round.

## References

1. Callaway, E. M. et al. A multimodal cell census and atlas of the mammalian primary motor cortex. Nature 598, 86–102 (2021).

2. Scala, F. et al. Phenotypic variation of transcriptomic cell types in mouse motor cortex. Nature 598, 144–150 (2021).

3. Tasic, B. et al. Adult mouse cortical cell taxonomy revealed by single cell transcriptomics. Nat. Neurosci. 19, 335–346 (2016).

4. Tasic, B. et al. Shared and distinct transcriptomic cell types across neocortical areas. Nature 563, 72–78 (2018).

5. Zeisel, A. et al. Brain structure. Cell types in the mouse cortex and hippocampus revealed by single-cell RNA-seq. Science 347, 1138–42 (2015).

6. Paul, A. et al. Transcriptional Architecture of Synaptic Communication Delineates GABAergic Neuron Identity. Cell 171, 522–539.e20 (2017).

7. Zeisel, A. et al. Molecular Architecture of the Mouse Nervous System. Cell 174, 999–1014.e22 (2018).

8. Gouwens, N. W. et al. Integrated Morphoelectric and Transcriptomic Classification of Cortical GABAergic Cells. Cell 183, 935–953.e19 (2020).

9. Cajal, S. R. y. Texture of the Nervous System of Man and the Vertebrates: Volume III An annotated and edited translation of the original Spanish text with the additions of the French version by Pedro Pasik and Tauba Pasik. (Springer-Verlag, 2002).

10. Lorente de Nó, R. La corteza cerebral del ratón. Trab. Lab. Invest. Bio.(Madrid) 20, 41–78 (1922).

11. Szentágothai, J. The Ferrier Lecture, 1977 The neuron network of the cerebral cortex: a functional interpretation. Proceedings of the Royal Society of London. Series B. Biological Sciences 201, 219–248 (1978).

12. Gokce, O. et al. Cellular Taxonomy of the Mouse Striatum as Revealed by Single-Cell RNA-Seq. Cell Rep 16, 1126–1137 (2016).

13. Harris, K. D. et al. Classes and continua of hippocampal CA1 inhibitory neurons revealed by single-cell transcriptomics. PLOS Biology 16, e2006387 (2018).

14. Klausberger, T. & Somogyi, P. Neuronal diversity and temporal dynamics: the unity of hippocampal circuit operations. Science 321, 53–7 (2008).

15. Abs, E. et al. Learning-Related Plasticity in Dendrite-Targeting Layer 1 Interneurons. Neuron 100, 684–699.e6 (2018).

16. Adesnik, H., Bruns, W., Taniguchi, H., Huang, Z. J. & Scanziani, M. A neural circuit for spatial summation in visual cortex. Nature 490, 226–31 (2012).

17. Cardin, J. A. et al. Driving fast-spiking cells induces gamma rhythm and controls sensory responses. Nature 459, 663–7 (2009).

18. Cardin, J. A. et al. Targeted optogenetic stimulation and recording of neurons in vivo using cell-type-specific expression of Chan-nelrhodopsin-2. Nat Protoc 5, 247–54 (2010).

19. Dipoppa, M. et al. Vision and Locomotion Shape the Interactions between Neuron Types in Mouse Visual Cortex. Neuron 98, 602–615.e8 (2018).

20. Dudok, B. et al. Alternating sources of perisomatic inhibition during behavior. Neuron 109, 997–1012.e9 (2021).

21. Fu, Y. et al. A cortical circuit for gain control by behavioral state. Cell 156, 1139–52 (2014).

22. Hofer, S. B. et al. Differential connectivity and response dynamics of excitatory and inhibitory neurons in visual cortex. Nat Neurosci 14, 1045–52 (2011).

23. Kamigaki, T. & Dan, Y. Delay Activity of Specific Prefrontal Interneuron Subtypes Modulates Memory-Guided Behavior. Nat Neurosci 20, 854–863 (2017).

24. Kvitsiani, D. et al. Distinct behavioural and network correlates of two interneuron types in prefrontal cortex. Nature 498, 363–366 (2013).

25. Lee, S., Kruglikov, I., Huang, Z. J., Fishell, G. & Rudy, B. A disinhibitory circuit mediates motor integration in the somatosensory cortex. Nature neuroscience 16, 1662–70 (2013).

26. Letzkus, J. J. et al. A disinhibitory microcircuit for associative fear learning in the auditory cortex. Nature 480, 331–5 (2011).

27. Lima, S. Q., Hromadka, T., Znamenskiy, P. & Zador, A. M. PINP: a new method of tagging neuronal populations for identification during in vivo electrophysiological recording. PLoS One 4, e6099 (2009).

28. Ma, W. et al. Visual Representations by Cortical Somatostatin Inhibitory Neurons—Selective But with Weak and Delayed Responses. J. Neurosci. 30, 14371–14379 (2010).

29. Pakan, J. M. et al. Behavioral-state modulation of inhibition is context-dependent and cell type specific in mouse visual cortex. eLife 5, e14985 (2016).

30. Pi, H. J. et al. Cortical interneurons that specialize in disinhibitory control. Nature 503, 521–4 (2013).

31. Pinto, L. & Dan, Y. Cell type-specific activity in prefrontal cortex during goal-directed behavior. Neuron 87, 437–450 (2015).

32. Polack, P. O., Friedman, J. & Golshani, P. Cellular mechanisms of brain state-dependent gain modulation in visual cortex. Nature neuroscience (2013) doi:10.1038/nn.3464.

33. Reimer, J. et al. Pupil fluctuations track fast switching of cortical states during quiet wakefulness. Neuron 84, 355–62 (2014).

34. Royer, S. et al. Control of timing, rate and bursts of hippocampal place cells by dendritic and somatic inhibition. Nature Neuroscience 15, 769–775 (2012).

35. Geiller, T. et al. Large-Scale 3D Two-Photon Imaging of Molecularly Identified CA1 Interneuron Dynamics in Behaving Mice. Neuron 108, 968–983.e9 (2020).

36. Kerlin, A. M., Andermann, M. L., Berezovskii, V. K. & Reid, R. C. Broadly tuned response properties of diverse inhibitory neuron subtypes in mouse visual cortex. Neuron 67, 858–71 (2010).

37. Khan, A. G. et al. Distinct learning-induced changes in stimulus selectivity and interactions of GABAergic interneuron classes in visual cortex. Nat Neurosci 21, 851–859 (2018).

38. Lovett-Barron, M. et al. Ancestral Circuits for the Coordinated Modulation of Brain State. Cell 171, 1411–1423.e17 (2017).

39. Lovett-Barron, M. et al. Multiple convergent hypothalamusbrainstem circuits drive defensive behavior. Nat Neurosci 23, 959–967 (2020).

40. Xu, S. et al. Behavioral state coding by molecularly defined paraventricular hypothalamic cell type ensembles. Science 370, eabb2494 (2020).

41. Qian, X. et al. Probabilistic cell typing enables fine mapping of closely related cell types in situ. Nature Methods 1–6 (2019) doi:10.1038/s41592-019-0631-4.

42. Chen, X. et al. High-Throughput Mapping of Long-Range Neuronal Projection Using In Situ Sequencing. Cell 179, 772–786.e19 (2019).

43. Ke, R. et al. In situ sequencing for RNA analysis in preserved tissue and cells. Nature methods 10, 857–60 (2013).

44. Nilsson, M. et al. Padlock Probes: Circularizing Oligonucleotides for Localized DNA Detection. Science 265, 2085–2088 (1994).

45. Sun, Y.-C. et al. Integrating barcoded neuroanatomy with spatial transcriptional profiling reveals cadherin correlates of projections shared across the cortex. bioRxiv 2020.08.25.266460 (2020) doi:10.1101/2020.08.25.266460.

46. McGinley, M. J., David, S. V. & McCormick, D. A. Cortical Membrane Potential Signature of Optimal States for Sensory Signal Detection. Neuron 87, 179–192 (2015).

47. McGinley, M. J. et al. Waking State: Rapid Variations Modulate Neural and Behavioral Responses. Neuron 87, 1143–1161 (2015).

48. Niell, C. M. & Stryker, M. P. Modulation of visual responses by behavioral state in mouse visual cortex. Neuron 65, 472–9 (2010).

49. Stringer, C. et al. Spontaneous behaviors drive multidimensional, brainwide activity. Science 364, 255 (2019).

50. Einstein, M. C., Polack, P.-O., Tran, D. T. & Golshani, P. Visually Evoked 3–5 Hz Membrane Potential Oscillations Reduce the Responsiveness of Visual Cortex Neurons in Awake Behaving Mice. J. Neurosci. 37, 5084–5098 (2017).

51. Jacobs, E. A. K., Steinmetz, N. A., Peters, A. J., Carandini, M. & Harris, K. D. Cortical State Fluctuations during Sensory Decision Making. Curr Biol 30, 4944–4955.e7 (2020).

52. Nestvogel, D. B. & McCormick, D. A. Visual thalamocortical mechanisms of waking state-dependent activity and alpha oscillations. Neuron 0, (2021).

53. Senzai, Y., Fernandez-Ruiz, A. & Buzsáki, G. Layer-Specific Physiological Features and Interlaminar Interactions in the Primary Visual Cortex of the Mouse. Neuron 101, 500–513.e5 (2019).

54. Schneider-Mizell, C. M. et al. Chandelier cell anatomy and function reveal a variably distributed but common signal. 2020.03.31.018952 https://www.biorxiv.org/content/10.1101/2020.03.31.018952v1 (2020) doi:10.1101/2020.03.31.018952.

55. Liu, B. H. et al. Visual receptive field structure of cortical inhibitory neurons revealed by two-photon imaging guided recording. The Journal of neuroscience: the official journal of the Society for Neuroscience 29, 10520–32 (2009).

56. Niell, C. M. & Stryker, M. P. Highly selective receptive fields in mouse visual cortex. The Journal of neuroscience: the official journal of the Society for Neuroscience 28, 7520–36 (2008).

57. Runyan, C. A. et al. Response features of parvalbumin-expressing interneurons suggest precise roles for subtypes of inhibition in visual cortex. Neuron 67, 847–57 (2010).

58. Buzsaki, G. et al. Nucleus basalis and thalamic control of neocortical activity in the freely moving rat. J Neurosci 8, 4007–26 (1988).

59. Castro-Alamancos, M. A. & Gulati, T. Neuromodulators Produce Distinct Activated States in Neocortex. J Neurosci 34, 12353–12367 (2014).

60. Eggermann, E., Kremer, Y., Crochet, S. & Petersen, C. C. H. Cholinergic signals in mouse barrel cortex during active whisker sensing. Cell Rep 9, 1654–1660 (2014).

61. Kalmbach, A. & Waters, J. Modulation of high- and low-frequency components of the cortical local field potential via nicotinic and muscarinic acetylcholine receptors in anesthetized mice. J Neurophysiol 111, 258–272 (2014).

62. Metherate, R., Cox, C. L. & Ashe, J. H. Cellular bases of neocortical activation: modulation of neural oscillations by the nucleus basalis and endogenous acetylcholine. J.Neurosci. 12, 4701–4711 (1992).

63. Pinto, L. et al. Fast modulation of visual perception by basal forebrain cholinergic neurons. Nature Neuroscience 16, 1857–1863 (2013).

64. Vanderwolf, C. H. An odyssey through the brain, behavior, and the mind. (Kluwer Academic Pub, 2003).

65. Arroyo, S., Bennett, C., Aziz, D., Brown, S. P. & Hestrin, S. Prolonged disynaptic inhibition in the cortex mediated by slow, non-alpha7 nicotinic excitation of a specific subset of cortical interneurons. The Journal of neuroscience: the official journal of the Society for Neuroscience 32, 3859–64 (2012).

66. Gasselin, C., Hohl, B., Vernet, A., Crochet, S. & Petersen, C. C. H. Cell-type-specific nicotinic input disinhibits mouse barrel cortex during active sensing. Neuron 109, 778–787.e3 (2021).

67. Gulledge, A. T., Park, S. B., Kawaguchi, Y. & Stuart, G. J. Heterogeneity of phasic cholinergic signaling in neocortical neurons. J Neurophysiol 97, 2215–2229 (2007).

68. Kawaguchi, Y. Selective cholinergic modulation of cortical GA-BAergic cell subtypes. J Neurophysiol 78, 1743–7 (1997).

69. Muñoz, W., Tremblay, R., Levenstein, D. & Rudy, B. Layer-specific modulation of neocortical dendritic inhibition during active wakefulness. Science 355, 954–959 (2017).

70. Xiang, Z., Huguenard, J. R. & Prince, D. A. Cholinergic switching within neocortical inhibitory networks. Science 281, 985–988 (1998).

71. Millman, D. J. et al. VIP interneurons in mouse primary visual cortex selectively enhance responses to weak but specific stimuli. eLife 9, e55130 (2020).

72. Cohen-Kashi Malina, K. et al. NDNF interneurons in layer 1 gain-modulate whole cortical columns according to an animal’s behavioral state. Neuron 109, 2150–2164.e5 (2021).

73. Pawelzik, H., Hughes, D. I. & Thomson, A. M. Physiological and morphological diversity of immunocytochemically defined parvalbumin- and cholecystokinin-positive interneurones in CA1 of the adult rat hippocampus. J. Comp. Neurol. 443, 346–367 (2002).

74. Gil, Z., Connors, B. W. & Amitai, Y. Differential regulation of neocortical synapses by neuromodulators and activity. Neuron 19, 679–86 (1997).

75. Harris, K. D. & Thiele, A. Cortical state and attention. Nat Rev Neurosci 12, 509–23 (2011).

76. Hasselmo, M. E. Neuromodulation and cortical function: modeling the physiological basis of behavior. Behav.Brain Res. 67, 1–27 (1995).

77. Kimura, F., Fukuda, M. & Tsumoto, T. Acetylcholine suppresses the spread of excitation in the visual cortex revealed by optical recording: possible differential effect depending on the source of input. European Journal of Neuroscience 11, 3597–3609 (1999).

78. Roberts, M. J. et al. Acetylcholine dynamically controls spatial integration in marmoset primary visual cortex. J Neurophysiol 93, 2062–72 (2005).

79. Bodor, A. L. et al. Endocannabinoid signaling in rat somatosensory cortex: laminar differences and involvement of specific interneuron types. J Neurosci 25, 6845–6856 (2005).

80. Harris, K. D. & Shepherd, G. M. The neocortical circuit: themes and variations. Nature neuroscience 18, 170–181 (2015).

81. Xu, H., Jeong, H. Y., Tremblay, R. & Rudy, B. Neocortical somatostatin-expressing GABAergic interneurons disinhibit the thalamorecipient layer 4. Neuron 77, 155–67 (2013).

## References

82. Wagor, E., Mangini, N. J. & Pearlman, A. L. Retinotopic organization of striate and extrastriate visual cortex in the mouse. Journal of Comparative Neurology 193, 187–202 (1980).

83. Pologruto, T. A., Sabatini, B. L. & Svoboda, K. ScanImage: Flexible software for operating laser scanning microscopes. BioMedical Engineering OnLine 2, 13 (2003).

84. Pachitariu, M. et al. Suite2p: beyond 10,000 neurons with standard two-photon microscopy. BioRxiv 061507, (2016).

85. Pachitariu, M., Stringer, C. & Harris, K. D. Robustness of Spike Deconvolution for Neuronal Calcium Imaging. J. Neurosci. 38, 7976–7985 (2018).

86. Chen, K. H., Boettiger, A. N., Moffitt, J. R., Wang, S. & Zhuang, X. RNA imaging. Spatially resolved, highly multiplexed RNA profiling in single cells. Science 348, aaa6090 (2015).

87. Moffitt, J. R. et al. Molecular, spatial, and functional single-cell profiling of the hypothalamic preoptic region. Science 362, (2018).

88. Lubeck, E., Coskun, A. F., Zhiyentayev, T., Ahmad, M. & Cai, L. Single-cell in situ RNA profiling by sequential hybridization. Nat Meth 11, 360–361 (2014).

89. Codeluppi, S. et al. Spatial organization of the somatosensory cortex revealed by osmFISH. Nat Methods 15, 932–935 (2018).

90. Eng, C.-H. L. et al. Transcriptome-scale super-resolved imaging in tissues by RNA seqFISH+. Nature 568, 235–239 (2019).

91. Lee, J. H. et al. Highly multiplexed subcellular RNA sequencing in situ. Science 343, 1360–3 (2014).

92. Chen, X., Sun, Y.-C., Church, G. M., Lee, J. H. & Zador, A. M. Efficient in situ barcode sequencing using padlock probe-based BaristaSeq. Nucleic Acids Res 46, e22–e22 (2018).

93. Wang, X. et al. Three-dimensional intact-tissue sequencing of single-cell transcriptional states. Science (2018) doi:10.1126/science.aat5691.

94. Sylwestrak, E. L., Rajasethupathy, P., Wright, M. A., Jaffe, A. & Deisseroth, K. Multiplexed Intact-Tissue Transcriptional Analysis at Cellular Resolution. Cell 164, 792–804 (2016).

95. Choi, H. M. T. et al. Third-generation in situ hybridization chain reaction: multiplexed, quantitative, sensitive, versatile, robust. Development 145, (2018).

96. Nagendran, M., Riordan, D. P., Harbury, P. B. & Desai, T. J. Automated cell-type classification in intact tissues by single-cell molecular profiling. eLife 7, e30510 (2018).

97. Kishi, J. Y. et al. SABER amplifies FISH: enhanced multiplexed imaging of RNA and DNA in cells and tissues. Nat. Methods 16, 533–544 (2019).

98. Ståhl, P. L. et al. Visualization and analysis of gene expression in tissue sections by spatial transcriptomics. Science 353, 78–82 (2016).

99. Vickovic, S. et al. High-definition spatial transcriptomics for in situ tissue profiling. Nat. Methods 16, 987–990 (2019).

100. Rodriques, S. G. et al. Slide-seq: A scalable technology for measuring genome-wide expression at high spatial resolution. Science 363, 1463–1467 (2019).

101. Chen, F. et al. Nanoscale Imaging of RNA with Expansion Microscopy. Nat Methods 13, 679–684 (2016).

102. Alon, S. et al. Expansion sequencing: Spatially precise in situ transcriptomics in intact biological systems. Science 371, eaax2656 (2021).

103. Xu, Q., Schlabach, M. R., Hannon, G. J. & Elledge, S. J. Design of 240,000 orthogonal 25mer DNA barcode probes. Proc Natl Acad Sci U S A 106, 2289–2294 (2009).

104. Reed, I. S. & Solomon, G. Polynomial Codes Over Certain Finite Fields. Journal of the Society for Industrial and Applied Mathematics 8, 300–304 (1960).

105. Moffitt, J. R. & Zhuang, X. Chapter One - RNA Imaging with Multiplexed Error-Robust Fluorescence In Situ Hybridization (MERFISH). in Methods in Enzymology (eds. Filonov, G. S. & Jaffrey, S. R.) vol. 572 1–49 (Academic Press, 2016).

106. Pertuz, S., Puig, D., Garcia, M. A. & Fusiello, A. Generation of All-in-Focus Images by Noise-Robust Selective Fusion of Limited Depth-of-Field Images. IEEE Transactions on Image Processing 22, 1242–1251 (2013).

107. Elad, M. Sparse and Redundant Representations: From Theory to Applications in Signal and Image Processing. (Springer-Verlag, 2010).

108. Viney, T. J. et al. Network state-dependent inhibition of identified hippocampal CA3 axo-axonic cells in vivo. Nat. Neurosci. 16, 1802–1811 (2013).

109. Xu, X., Roby, K. D. & Callaway, E. M. Mouse cortical inhibitory neuron type that coexpresses somatostatin and calretinin. J. Comp. Neurol. 499, 144–160 (2006).

110. Xu, X., Roby, K. D. & Callaway, E. M. Immunochemical characterization of inhibitory mouse cortical neurons: Three chemically distinct classes of inhibitory cells. J Comp Neurol 518, 389–404 (2010).

111. Schuman, B. et al. Four Unique Interneuron Populations Reside in Neocortical Layer 1. J Neurosci 39, 125–139 (2019).

112. Gesuita, L. & Karayannis, T. A ‘Marginal’ tale: the development of the neocortical layer 1. Curr Opin Neurobiol 66, 37–47 (2021).

113. Kawaguchi, Y. & Kubota, Y. GABAergic cell subtypes and their synaptic connections in rat frontal cortex. Cereb.Cortex 7, 476–486 (1997).

114. Kubota, Y. & Kawaguchi, Y. Two distinct subgroups of cholecystokinin-immunoreactive cortical interneurons. Brain Res 752, 175–183 (1997).

115. Harvey, C. D., Collman, F., Dombeck, D. A. & Tank, D. W. Intracellular dynamics of hippocampal place cells during virtual navigation. Nature 461, 941–6 (2009).

116. Stringer, C., Pachitariu, M., Steinmetz, N., Carandini, M. & Harris, K. D. High-dimensional geometry of population responses in visual cortex. Nature 571, 361–365 (2019).

117. Meehan, C., Meehan, S. & Moore, W. Uniform Manifold Approximation and Projection (UMAP). MATLAB Central File Exchange Available at: https://www.mathworks.com/matlabcentral/fileexchange/71902 (2020).

